# Simple visualization of submicroscopic protein clusters with a phase-separation-based fluorescent reporter

**DOI:** 10.1101/2022.07.13.499962

**Authors:** Thomas R. Mumford, Diarmid Rae, Emily Brackhahn, Abbas Idris, David Gonzalez-Martinez, Ayush Aditya Pal, Michael C. Chung, Juan Guan, Elizabeth Rhoades, Lukasz J. Bugaj

**Affiliations:** Department of Bioengineering, University of Pennsylvania, Philadelphia, PA, 19104, USA; Department of Chemistry, University of Pennsylvania, Philadelphia, PA, 19104, USA; Department of Physics, University of Florida, Gainesville, FL 32611; Department of Anatomy and Cell Biology, University of Florida, Gainesville, FL 32611; Biochemistry and Molecular Biophysics Graduate Group, Perelman School of Medicine, University of Pennsylvania, Philadelphia, PA, 19104, USA; Abramson Cancer Center, University of Pennsylvania, Philadelphia, PA, 19104, USA; Institute of Regenerative Medicine, University of Pennsylvania, Philadelphia, PA, 19104, USA

## Abstract

Protein clustering plays numerous roles in cell physiology and disease. However, protein oligomers can be difficult to detect because they are often too small to appear as puncta in conventional fluorescence microscopy. Here we describe a fluorescent reporter strategy that detects protein clusters with high sensitivity, called CluMPS (<u>Clu</u>sters <u>M</u>agnified by <u>P</u>hase <u>S</u>eparation). A CluMPS reporter detects and visually amplifies even small clusters of a binding partner, generating large, quantifiable fluorescence condensates. We use computational modeling and optogenetic clustering to demonstrate that CluMPS can detect small oligomers and behaves rationally according to key system parameters. CluMPS detected small aggregates of pathological proteins where the corresponding GFP fusions appeared diffuse. CluMPS also detected and tracked clusters of unmodified and tagged endogenous proteins, and orthogonal CluMPS probes could be multiplexed in cells. CluMPS provides a powerful yet straightforward approach to observe higher-order protein assembly in its native cellular context.

## Introduction

Protein clustering is ubiquitous and plays important roles throughout normal and disease physiology^1–4^. Processes including transmembrane receptor signaling, transcription factor activation, and cellular stress responses all depend on the appropriate formation of protein clusters^5–9^. Aberrant clustering, in turn, has been linked with numerous pathologies, including in neurodegenerative diseases, cancer, and aging^11,14,16,18,20,22,24,26,71,74^.

Clustering is driven by multivalent interactions whose characteristics generate diverse biophysical, biochemical, and functional outcomes^3,10,12^. Protein assemblies can serve many functions, including enhancing biochemical reaction rates, sequestering proteins, spatially restricting biochemical reactions, and buffering concentrations of cellular components^4,12,13,15^. Such assemblies can also play roles in cell signal processing, enabling ultra-sensitivity, bistability, and memory^2,17,19^. A deeper understanding of when/where protein clustering occurs will not only paint a clearer picture of cellular organization and function but will also provide novel control nodes where clustering can be modulated to regulate cell behavior for biomedicine and biotechnology.

It is currently challenging to understand the extent of protein clustering in cells because we lack straightforward tools to observe it. Physiologically relevant clusters are often too small to see using light microscopy^21,23,25,27,28^. The toxic “soluble oligomers’’ of α-synuclein, amyloid-β, and TDP-43 found in neurodegenerative diseases consist of only 10s of monomers, in assemblies roughly 10-100 nm in size^14,16,39,42,71,72^. In signaling, protein nanoclusters transduce signals from activated T cell receptors^31^, and ∼100 nm clusters of scaffold proteins compartmentalize protein kinase A activity^32^. Moreover, it is estimated that most proteins form homo-oligomers^33^, and that at least 18% of the human proteome is organized into condensates > ∼100 nm in size^34^. Direct observation of such small clusters is often not possible with simple fluorescent protein fusions because the clusters do not concentrate enough fluorophores to exceed the fluorescence background. This principle is demonstrated by *A. thaliana* Cryptochrome 2 (Cry2), which clusters upon blue light stimulation^35^. Cry2 cluster size is a function of its concentration, and below a certain concentration, clusters of fluorescent fusions of Cry2 do not appear^35,50^. Nevertheless, small clusters continue to form in this low concentration regime, evidenced by the fact that certain optogenetic tools based on Cry2 clustering can still be activated despite the absence of visible clusters^34,53^. Additionally, large Cry2 clusters can be generated at low concentrations by co-expression with multivalent binding partners^38–47^ or by appending Cry2 with certain peptides or disordered regions^38,40,41,43–49,50^, suggesting that small clusters are forming in the absence of these additional domains.

Specialized microscopy methods exist to observe such small clusters, including Foerster resonance energy transfer (FRET), fluorescence anisotropy, fluorescence correlation spectroscopy, and super-resolution imaging. However, these methods are subject to drawbacks including complexity in acquisition and analysis, low signal-to-noise ratio, significant optimization for each target, long acquisition times that obscure fast dynamic changes, or the need for protein overexpression^51^. Overexpression can yield artifacts because clustering is non-linearly dependent on concentration, thus raising questions of whether the observed clustering is representative of the native cellular context^67,69^.

We sought to address this challenge with a fluorescent reporter of protein clustering that 1) could detect small clusters, 2) is simple to design and image, 3) is easily adaptable to new targets, 4) could report on endogenous targets, and 5) could be rapidly imaged in live mammalian cells. Here we introduce such a reporter strategy called CluMPS (<u>Clu</u>sters <u>M</u>agnified via <u>P</u>hase <u>S</u>eparation). CluMPS is a multivalent fluorescent reporter that detects and amplifies clusters of a target protein, producing large phase-separated condensates that can be easily measured. Combining live cell imaging, optogenetic clustering, and stochastic modeling, we validated the CluMPS approach, demonstrated its sensitivity to small clusters, and determined critical system features required for CluMPS activation. We then demonstrated that CluMPS can visualize diverse pathological clusters that otherwise appear diffuse, including those of endogenous proteins, can report on drug-induced cluster dynamics, and can be multiplexed to visualize distinct clustered species in the same cell. Our work thus provides a modular reporter strategy for sensitive and dynamic observation of protein clusters in mammalian cells.

## Results

### Design of the CluMPS reporter

The CluMPS reporter leverages phase separation to detect and visually amplify small protein clusters. We chose phase separation as a readout due to its sensitivity and simplicity: phase separation can arise from multivalent interactions of even trimers and tetramers^69^, resulting in micron-scale condensates that are easy to detect.

A CluMPS reporter is multivalent and has affinity for a target protein of interest. If the target is clustered, CluMPS and the target condense together into large fluorescent puncta. Otherwise, CluMPS remains diffuse (**Figure 1A**). Such conditional condensation of CluMPS could be triggered by even small oligomers of its target, allowing CluMPS to report on otherwise invisible protein clusters. CluMPS thus would provide a straightforward and modular method to detect submicroscopic clustering in live cells and could be readily adapted to report on endogenous proteins, thus fulfilling our design criteria.

**Figure 1:**
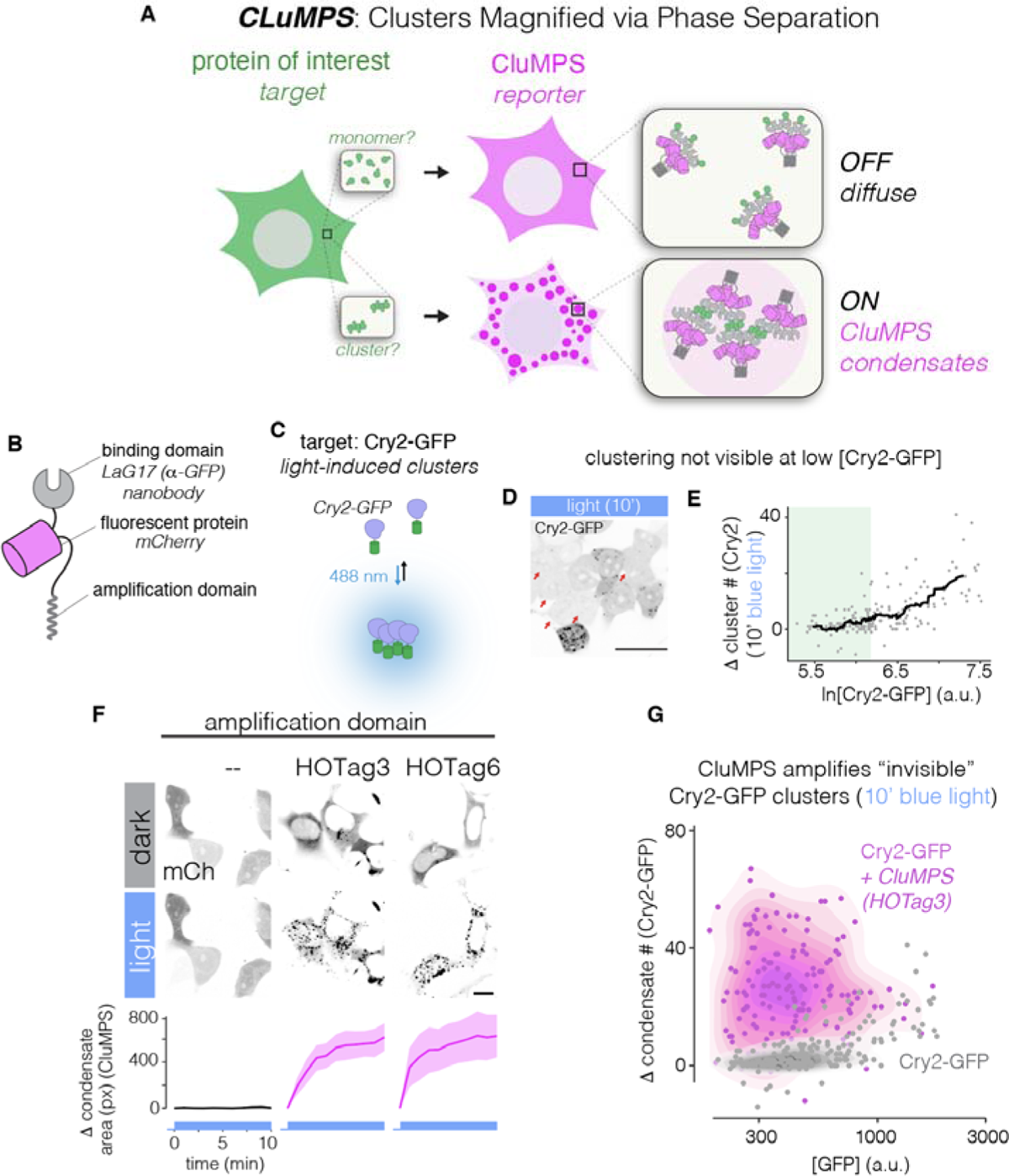
CluMPS reporters amplify small protein oligomers. **(A)** A CluMPS reporter (magenta) forms large condensates when its target (green) is clustered but remains diffuse when the target is monomeric. **(B)** A CluMPS reporter consists of a target binding domain, fluorescent protein, and an amplification domain that facilitates phase separation in the presence of a clustered target. **(C)** Cry2-GFP is diffuse in the dark but clusters when stimulated with blue light. **(D)** Clusters of Cry2-GFP can be seen in high-expressing cells after 10 minutes of stimulation with blue light, but low-expressing cells (red arrows) do not show visible clusters. Scale bars = 20 µm. **(E)** Cry2-GFP clustering is concentration dependent, and at low concentrations (green), most light-induced Cry2-GFP clusters are too small to see. **(F)** CluMPS variants with HOTag3 or HOTag6 amplification domains successfully generated large phase-separated condensates upon optogenetic clustering of Cry2-GFP, which was expressed at levels where clusters are not visible in the absence of CluMPS. A control reporter construct without an amplification domain (left) shows no condensation. Data represent the means, error bars = 95% CI of approximately 20-50 cells per group. See also (**Supplementary** Figure 1). Scale bar = 10 µm. **(G)** Quantification of light-induced Cry2-GFP clusters, in the presence or absence of Lag17-CluMPS (HOTag3) shows CluMPS-induced amplification of Cry2-GFP clustering including at low concentrations of Cry2-GFP where, in the absence of CluMPS, clustering is undetectable.

A CluMPS reporter consists of three domains: 1) a binding domain for a target protein, 2) a fluorescent protein, and 3) an amplification domain that provides multivalency (**Figure 1B**). We tested five candidate amplification domains — two multimeric coiled-coils (hexameric HOTag3 and tetrameric HOTag6)^57^ and three intrinsically disordered regions (IDR: FUS(LC), LAF-1 RGG, Xvelo)^57^ — and we assayed their ability to enable amplification of small clusters of GFP. To target GFP, we used the GFP-binding nanobody LaG17 as the binding domain^58^.

To benchmark the performance of our candidate CluMPS designs, we used an optogenetic approach to induce aggregation of GFP. We fused GFP to Cry2, which forms clusters under blue light stimulation^37,52,59^(**Figure 1C**). The size of light-induced Cry2 clusters depends on concentration^36,50,77^. While Cry2-GFP forms large, visible clusters under high expression, clusters in low-expressing cells are too small to observe above the diffuse fluorescence background (**Figure 1D,E**). Such small unobservable clusters provided an ideal test-case for CluMPS amplification.

We tested the ability of candidate CluMPS constructs to form protein condensates when co-expressed with Cry2-GFP and stimulated with blue light. When expressed alone or with Cry2-GFP in the dark, all CluMPS variants showed diffuse expression (**Figure 1F, top panels, Supplementary** Figure 1A-C**, Supplementary Video 1, Supplementary** Figure 2). Upon light stimulation, CluMPS variants with disordered amplification domains showed little or no phase separation, comparable to the negative control (LaG17-mCh with no amplification domain). By contrast, constructs with multimeric amplification domains (HOTag3 or HOTag6) formed large condensates within 10s of seconds of stimulation (**Figure 1F, Supplementary** Figure 1A-C**, Supplementary Video 1**). CluMPS condensation could be observed at all concentrations of Cry2-GFP, including at low levels (**Figure 1G**). Thus, we identified at least two distinct amplification domains that could be used to implement the CluMPS approach. We used HOTag3 (6-mer) as the amplification domain for subsequent experiments, unless otherwise noted.

To determine if CluMPS could amplify other multivalent species, we co-expressed the LaG17-mCh-HOTag3 construct (LaG17-CluMPS) with a concatamer of five tandem GFP molecules (GFPx5), which appeared diffuse when expressed alone (**Supplementary** Figure 3A,B). CluMPS showed robust condensation in cells where it was co-expressed with GFPx5. (**Supplementary** Figure 3C**, D**). GFP concatamers were pentameric in cells and were not spontaneously forming higher order assemblies, as validated through in-cell fluorescence correlation spectroscopy (**Supplementary** Figure 4). To demonstrate CluMPS against a non-GFP target, we tested its activity against concatamers of the small ALFA peptide tag ^60^. We replaced LaG17 with a nanobody against ALFA (nbALFA-CluMPS), and we concatamerized four ALFA tags on a single peptide chain as the multivalent target (ALFAx4) (**Supplementary** Figure 3E). NbALFA-CluMPS was diffuse when expressed alone but formed micron-scale condensates when co-expressed with ALFAx4 (**Supplementary** Figure 3F-H). Thus CluMPS is a generalizable approach to detect multivalency in even small assemblies of proteins.

We considered whether CluMPS could report false positives, for example by phase separating with a target that is not constitutively multivalent but had a weak intrinsic propensity to self-associate. To test this possibility, we observed whether LaG17-CluMPS would phase separate with a fusion of GFP with either FUS(LC) or RGG, two disordered domains that are not multimeric but that have sufficient self-affinity to drive phase separation at high concentration or when multimerized^56^. In both cases, CluMPS did not induce phase separation at any expression level of the GFP fusions (**Supplementary** Figure 5), Thus, while CluMPS is activated by small clusters, we do not observe activation by monomers, even those with a latent ability to multimerize.

We also tested whether CluMPS could report on the dynamics of clustering by measuring CluMPS reversion to a diffuse state upon de-clustering of its target. We examined OFF-kinetics of CluMPS upon light removal and declustering of Cry2 and another light-induced clustering protein, BcLOV4^57^. In both cases, CluMPS reverted back to a diffuse state with a half-life of either ∼11 min (Cry2 target) or ∼100 s (BcLOV4 target), reflecting known differences in the OFF-kinetics of these targets^61,62^. Thus, CluMPS does not lock targets into condensates and can dynamically report on their clustered state (**Supplementary** Figure 6).

### A stochastic model reveals important parameters for CluMPS activation

To understand the key principles that govern CluMPS, we built an *in silico* model to rapidly explore the experimental parameter space. We extended a kinetic Monte Carlo model of protein clustering^63^ to simulate interactions between two multivalent species, a ‘target’ and a ‘reporter’ (CluMPS) (**Figure 2A**). Each species moved freely within its 2D grid but could interact with members of the opposite species near its corresponding grid position (**Figure 2B**). The valency of each unit defined the maximum number of such cross-species interactions per unit. A binding energy ΔE defined the strength of a single target:reporter interaction. Reporter units had a valency of six to approximate the valency of CluMPS probes containing HOTag3. During each simulation step, one target or reporter unit could move to an adjacent grid space. The likelihood of movement *p_move_* was determined by the enthalpy of cross-plane interactions, a function of ΔE and the number and valency of current binding partners (**Figure 2B**, see **Methods** for more details). Using this framework, we simulated through time the spatial distribution of targets and reporters as a function of four system parameters: 1) binding energy (ΔE), 2) target cluster size, 3) the fraction of target that was in a cluster, since proteins clusters can exist in a distribution of sizes within the cell, and 4) the concentrations of both target and reporter (**Figure 2C**). We began each simulation with randomly placed target and reporter units and ran simulations until a steady state of reporter condensates was reached. We then quantified the fraction of reporter units that were part of a cluster at steady state. Simulations of these simple rules of motion and interaction yielded large target/reporter condensates over time (**Figure 2D, Supplementary Video 2**). Both higher ΔE and higher target valency drove stronger condensation, recapitulating intuitive aspects of the CluMPS system (**Figure 2D,E**). Condensation increased sigmoidally with increasing binding affinity, with a more switch-like relationship at higher target valencies (**Figure 2E)**. Condensation was also sensitive to the fraction of target that was in a cluster (multivalent). We modeled cluster size distributions by adding monomeric target units until a predefined percentage of clustered targets was reached (from 25% to 100%). Increasing the amount of target monomers (decreasing the percentage of target clusters) diminished condensation because monomers that bound reporters lowered the likelihood of multivalent-multivalent interactions required for condensation (**Figure 2F**). Our simulations also suggested that the CluMPS strategy could robustly amplify clusters as small as tetramers, and, under ideal conditions (high affinity, high percent clustered) even dimers and trimers, though to a lesser extent (**Figure 2E,F**). Finally, our simulations suggested that CluMPS condensation would be sensitive to the relative concentrations of target and reporter, which we simulated by varying the number of target and reporter units in the simulation. Condensation was most robust when target and reporter were present in roughly equal amounts but was disfavored when either was in large excess (**Figure 2G)**. Additionally, simulations with high concentrations of target and reporter yielded condensation that was less sensitive to target:reporter affinity and target cluster size, given appropriate target:reporter ratio (**Figure 2H**). Collectively, these simulations are consistent with the established role of multivalency in driving phase separation and provide predictions for how to optimize and interpret CluMPS behavior^51,69,82^.

**Figure 2.**
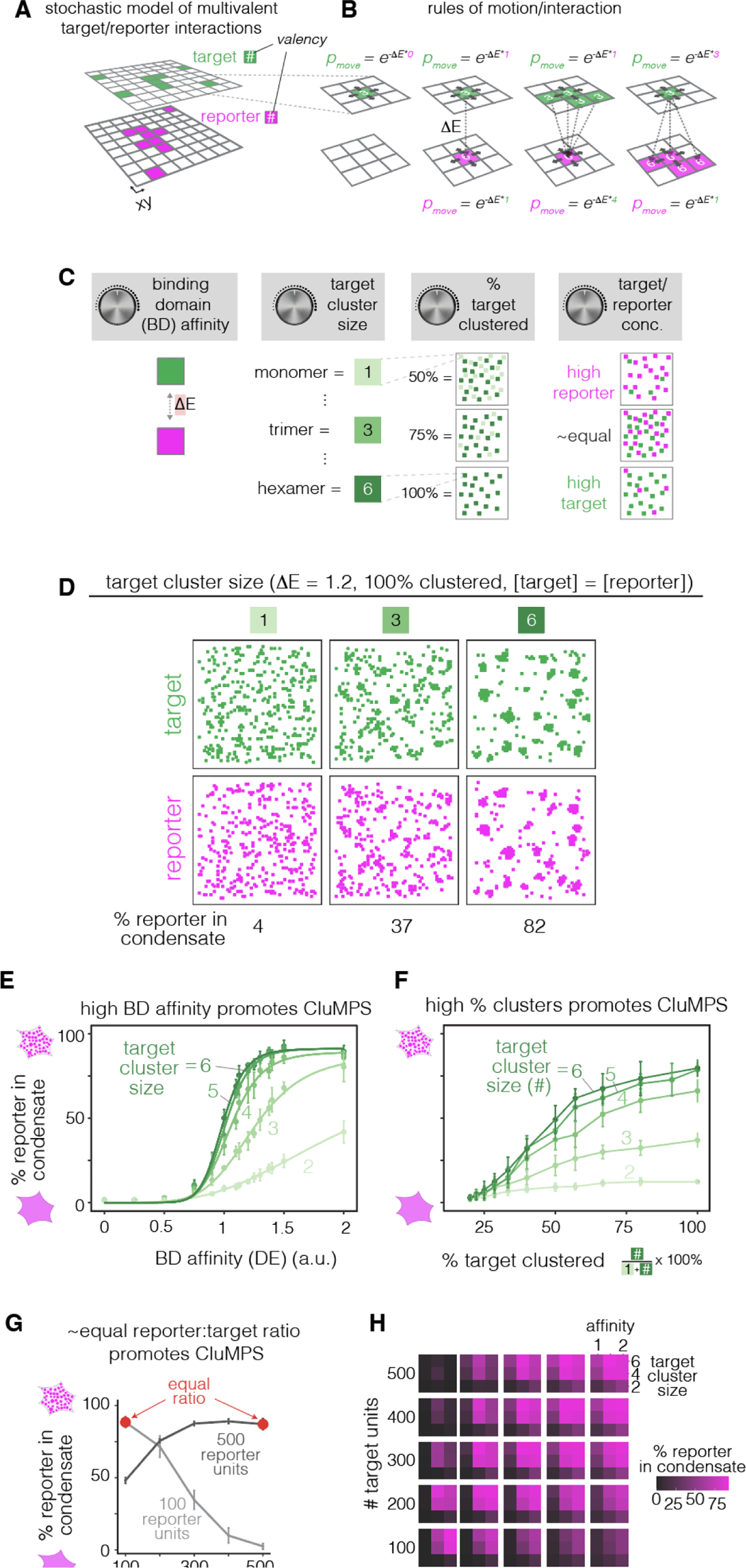
Stochastic model of CluMPS condensation. **(A)** Schematic of a kinetic Monte Carlo model of CluMPS/target interaction. CluMPS units (magenta) and target units (green) populate separate but interacting 2D grid planes. Both CluMPS and target units have a predetermined valency, which sets the number of adjacent units on the opposite plane with which they can interact. **(B)** During each simulation step a random unit is selected to move. The probability that the move is accepted is set by the enthalpy of cross-plane interactions of the unit’s initial state. The number of interactions is limited by the unit’s valency; i.e. if a unit set to move has four cross-plane neighbors, but a valency of three, the enthalpy will be calculated as if there are only three neighbors (right-most panel). See **Methods** for more details. **(C)** The model was used to understand how CluMPS parameters (binding affinity ΔE, target cluster size, percent target clustered, and concentration) influenced condensation. **(D)** Representative simulation results for different target valencies with constant binding energy and 100% target clustered. Simulated CluMPS/target condensates form more readily with larger target cluster sizes. See also **Supplementary Movie 2**. **(E)** Quantitation of simulations that sampled different binding domain affinities across target cluster sizes. Condensation increased with higher binding energy and larger target cluster sizes. Data points represent means ± 1 s.d. of 10 simulations. Trend lines are sigmoidal curves of best fit for each target cluster size. **(F)** Quantitation of simulations that sampled different distributions of target clustering and target cluster sizes. Addition of monovalent target suppresses CluMPS:target condensation for all target cluster sizes. Data points represent means ± 1 s.d. of 10 simulations. **(G)** The effect of CluMPS and target concentrations was tested by varying the number of units of each type. In simulations with target valency of 4 and target-reporter affinity of 2, increasing number of target units from 100-500 increased condensation when 500 reporter units were present, but decreased condensation when 100 reporter units were present. Data points represent the mean percent of reporter units in condensates of 10 simulations ± 1 s.d. **(H)** Heat map shows percent of reporter in condensates over all simulation parameters tested. Large 5×5 grid varies the number of target and reporter units used in each simulation, while each small 3×3 grid varies the affinity between target and reporter and the valency of target units. Color values were generated from the mean percent of reporter units in condensates for 10 simulations at each condition.

### Optogenetic test systems validate model predictions

We next sought to experimentally test our model predictions using two optogenetic clustering systems to generate target clusters (**Figure 3A**). To test the role of affinity, we measured amplification of Cry2-GFP using CluMPS reporters with distinct GFP-binding nanobodies of different affinities: LaG17-CluMPS (K_d_ = 50 nM), LaG6-CluMPS (310 nM), and LaG18-CluMPS (3800 nM) (**Figure 3B, Supplementary** Figure 7)^64–66,68,70^. In agreement with predictions, LaG17-CluMPS robustly amplified light-induced Cry2-GFP clusters, whereas the lower-affinity LaG6-CluMPS and LaG18-CluMPS were progressively less effective, forming smaller and fewer clusters after light stimulation (**Figure 3C, Supplementary** Figure 7**, Supplementary Video 3)**.

**Figure 3.**
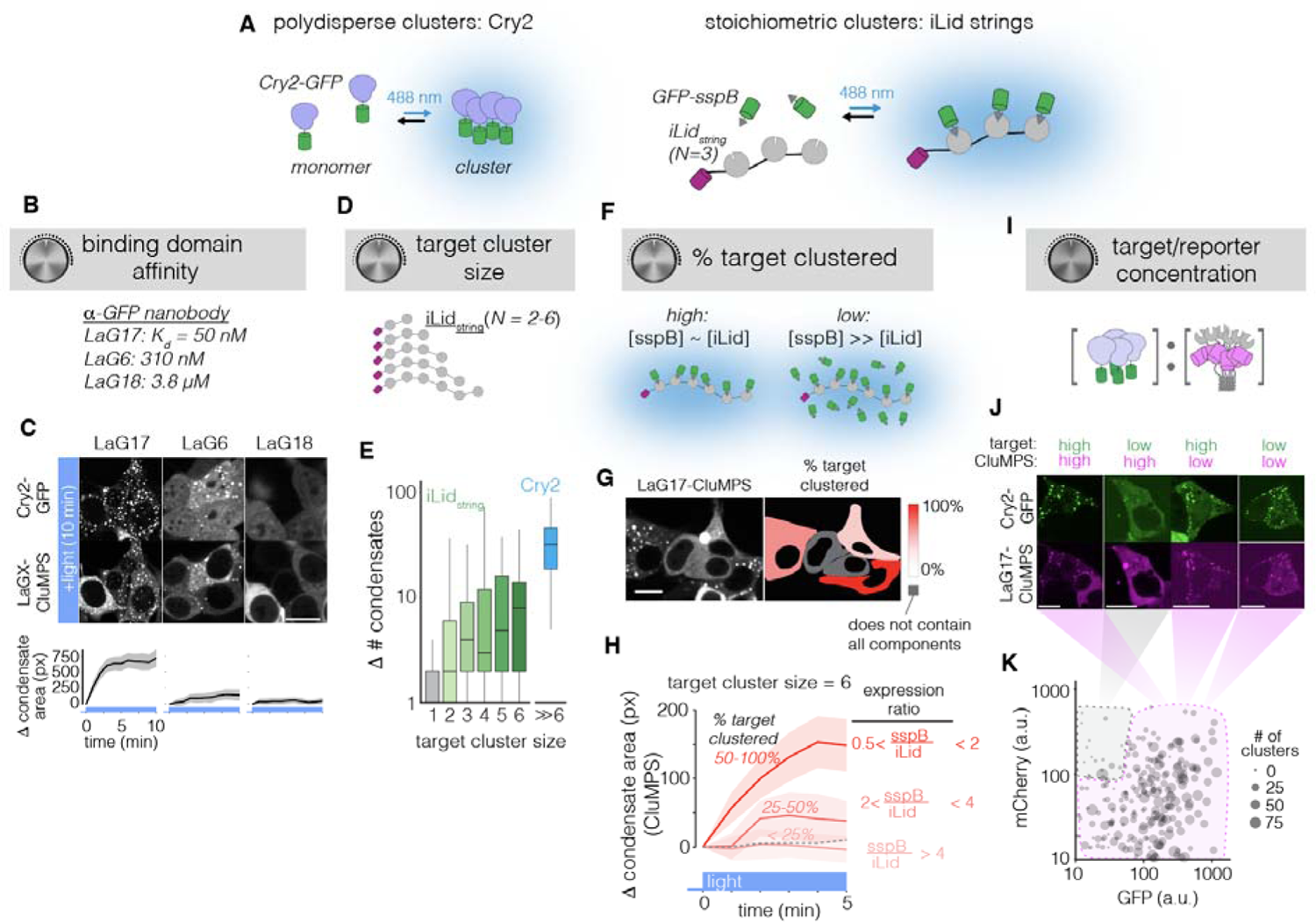
CluMPS activation is determined by binding affinity, target cluster size, cluster size distribution, and concentration. **(A)** Two optogenetic test systems were employed to empirically test the effects of cluster parameters on CluMPS activation. Cry2-GFP was used to generate large clusters of variable size. The iLid strings system was constructed to generate stoichiometrically-defined clusters. Multiple iLids were fused on a single peptide (iLid_string_), while sspB, which binds iLid in response to blue light, was monomeric and fused to GFP. (**B)** Amplification of Cry2-GFP clusters using CluMPS variants with different affinities of GFP-binding nanobody **(C)** LaG17 (K_d_ = 50 nM, left) yielded robust amplification, LaG6 (K_d_ = 310 nM, center) showed weaker condensation, and LaG18 (K_d_ = 3800 nM, right) did not produce condensation. Traces represent mean. Ribbons = 95% CI of approximately 100-200 cells per group. Scale bar = 20 µm. **(D)** iLid strings of sizes 2-6. **(E)** CluMPS activation in response to light-induced clustering of sspB-GFP with different iLid string sizes. CluMPS condensation increases with valency of the iLid string; all strings produce fewer condensates than Cry2-GFP. Data represents medians and quartiles of 80-200 cells per group. **(F)** The percentage of light-induced clusters in a given cell was estimated by examining the relative amounts of iLid string and GFP-sspB (see also **Supplementary** Figure 10). **(G)** Representative image of sspB-GFP/CluMPS condensation in cells that have iLidx6, GFP-sspB, and LaG17-CluMPS and were stimulated with blue light for five minutes. Color coding indicates the estimated percentage of GFP-sspB per cell that is clustered. Scale bar = 20 µm. (**H**) Quantification of relationship between light-induced CluMPS activation and the percentage of clustered target. Robust formation of GFP-sspB/CluMPS condensates is observed when 50-100% of sspB-GFP is clustered. Condensation is diminished when only 25-50% of the target is clustered, and is virtually undetectable when only < 25% of target is clustered. Dashed gray line represents negative control of cells with CluMPS and GFP-sspB but no iLid string. Traces represent means. Error bars = 95% CI of approximately 200 cells for the ‘50-100% clustered’ group and 40-60 cells for other groups. **(I)** Cry2-GFP with LaG17-CluMPS was used to understand how relative concentration of target and CluMPS affect phase separation upon blue-light induced multimerization of Cry2-GFP. **(J)** Representative images of CluMPS and Cry2-GFP in four expression regimes (high target + high clumps, low target + high clumps, high target + low CluMPS, and low target + low CluMPS). Clusters appeared in all regimes except low target + high CluMPS. Scale bars = 20 µm. **(K)** GFP (Cry2-GFP) and mCh (LaG17-mCh-HOTag3) intensities of single cells transfected with both Cry2-GFP and CluMPS plotted with dot size scaled to number of CluMPS condensates observed after 10 minute blue light stimulation. For similar analysis with GFPx5 and ALFAx4, see **Supplementary** Figures 13,14)

To test the roles of target cluster size and distribution, we adopted a method to generate stoichiometrically defined GFP clusters by concatemerizing the iLid protein (2-6mer “strings”, iLidx2-iLidx6), which forms heterodimers with sspB under blue light^69–71^(Figure 3A, **right**). In this system, blue light transitioned GFP-sspB from a monomer to a cluster, with cluster size defined by the string length. Co-transfection of the iRFP-iLid_string_, GFP-sspB, and LaG17-CluMPS allowed us to observe light-induced condensate formation with quantitative measures of expression of each component (**Supplementary** Figure 8).

We used this system to confirm that cluster size, modeled by iLid_string_ length, correlated with the magnitude of CluMPS amplification, as predicted by simulations (**Figure 3D,E**). Robust condensation was observed in the presence of iLidx6 and was progressively weaker with smaller string lengths (**Figure 3E, Supplementary** Figure 8D). The strong clustering in cells with iLidx6 was still lower compared to that observed in cells expressing Cry2-GFP, which forms clusters with valencies >> 6. In some cells, condensation could be observed with string sizes as small as 2 and 3, as predicted by our simulations. To further confirm these results, we co-expressed LaG17-CluMPS with tandem GFPs (2X or 3X) on a single peptide chain, and we observed robust clustering in most cells (**Supplementary** Figure 9). These results confirm that, under ideal conditions (stable multimers, high CluMPS:target affinity, 100% multimerized), CluMPS can report on target clusters as small as trimers and dimers. However, we expect that most ‘real-world’ conditions will be less ideal, and that such small oligomers will be challenging to detect.

We assessed the role of cluster size distribution using the iLid strings system by taking advantage of cell-to-cell variability in expression levels of each component. We estimated the percentage of GFP-sspB in iLid_string_-bound clusters by comparing the relative expression levels of iRFP-iLid_string_ and GFP-sspB (**Figure 3F, Supplementary** Figure 10). For example, a high sspB/iLid_string_ ratio resulted in a low percentage of GFP-sspB (the CluMPS target) clustered since there remained an excess of monomeric, unbound GFP-sspB even after light stimulation (**Figure 3F,G, Supplementary** Figure 10). We poly-transfected GFP-sspB and iLid_string_ plasmids to achieve uncorrelated expression and a wide range of sspB:iLid string ratios in a single pool of cells^71–73^ (**Supplementary** Figure 11). In line with simulations, we found that cells with a high percentage of clustered target (50-100%, 0.5 < sspB/iLid_string_ < 2) showed strong light-induced condensation, whereas cells with 25-50% (2 ≤ sspB/iLid_string_ < 4) showed less condensation, and at < 25% (sspB/iLid_string_ ≥ 4), condensation was not detectable (**Figure 3H, Supplementary** Figure 12).

Finally, we examined how CluMPS condensation was influenced by expression levels of target and reporter. We assayed a wide range of expression of LaG17-CluMPS and target for both Cry2-GFP and GFPx5 (**Figure 3I,J**, **Supplementary** Figure 13). For light activated Cry2-GFP, robust clustering was observed across expression levels, except in the case of ‘low’ Cry2-GFP and ‘high’ CluMPS (**Figure 3J,K, Supplementary** Figure 13C,D). Additionally, condensation decreased as the CluMPS:target ratio grew, as predicted by our simulations. Condensation was more sensitive to concentrations with the GFPx5 target, which has lower valency than light-activated Cry2-GFP and also does not self-associate. Here, robust condensation required sufficient expression of both components (**Supplementary** Figure 13E,F). Condensation was strongest at an intermediate CluMPS:GFPx5 ratio, with weaker condensation when relative expression was biased too far towards either component (**Supplementary** Figure 13G).

To obtain a conservative estimate of the lower limit of target concentration required to trigger CluMPS, we co-expressed a small target cluster (iRFP-ALFAx4) with nbALFA-GFP-HOTag3, and we measured condensation as a function of target and reporter fluorescence that was calibrated against know protein concentrations (**Supplementary** Figure 14A-E). Condensates formed below the lower limit of imaging detection of iRFP-ALFAx4, ∼800 nM, confirming that CluMPS can operate at concentrations typical of endogenous proteins (**Supplementary** Figure 14D). We emphasize, however, that the limit of detection will vary as a function of cluster size, distribution, target affinity, and ratio of target to reporter, as detailed above.

### CluMPS amplifies small clusters of pathological proteins

To determine whether CluMPS enhances visualization of known protein oligomers, we tested whether it could detect and amplify small aggregates of disease-associated proteins. These target proteins were fused with GFP to allow amplification with LaG17-CluMPS (LaG17 binds GFP with K_d_ = 50 nM).

The nucleocapsid (N) protein of SARS-CoV-2 can phase separate due to its self-affinity, disordered regions, and RNA interactions^74^. However, a GFP-N fusion appeared diffuse in HEK 293T cells, as previously observed^74^, likely because N-protein condensates are small (**Figure 4A**). By contrast, co-expression with LaG17-CluMPS resulted in micron-scale aggregates of both GFP-N and the reporter, observed in 100% of cells (**Figure 4A-C**).

**Figure 4.**
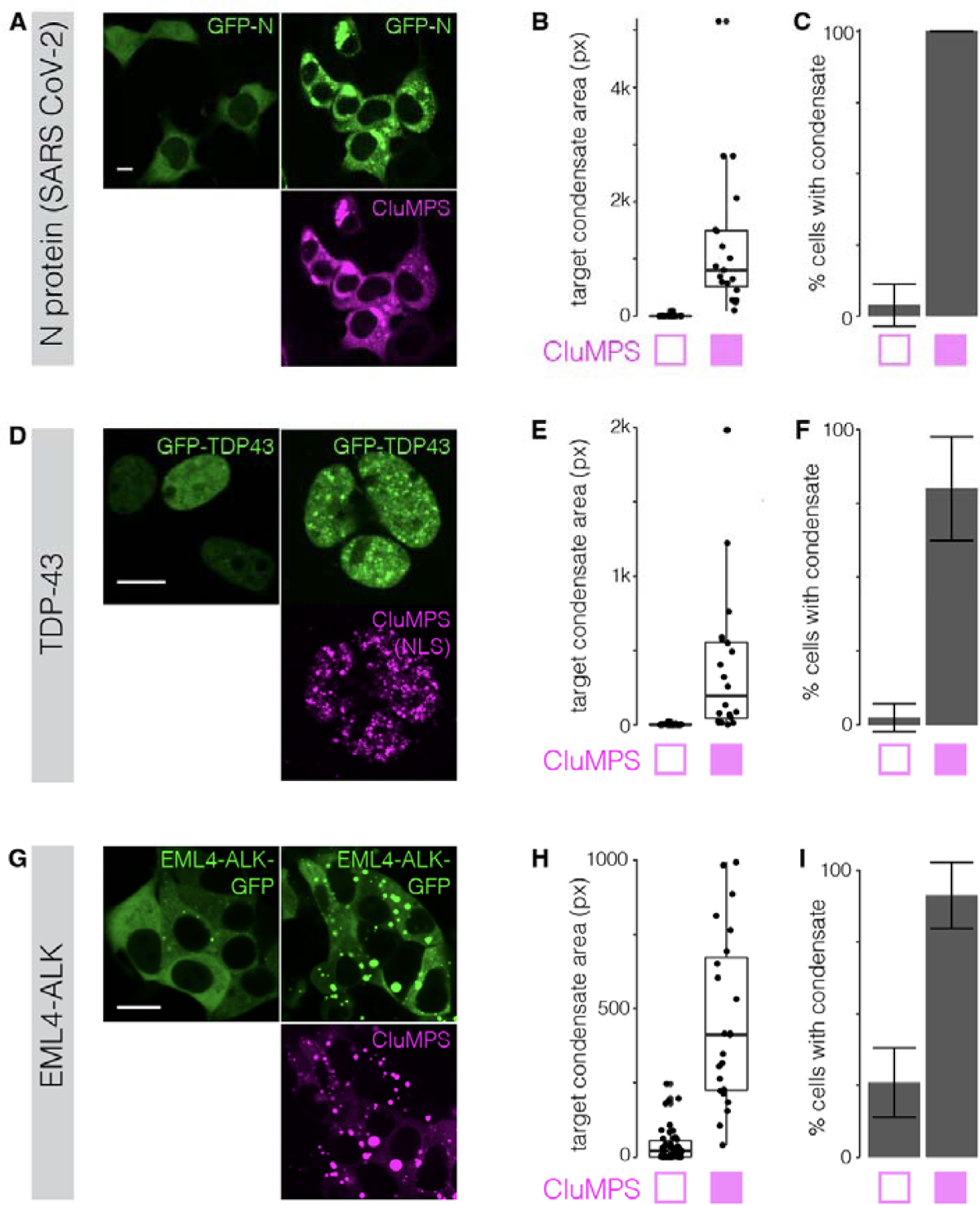
CluMPS detects clustering of pathological proteins. **(A)** SARS-CoV-2 nucleocapsid protein (N) with N-terminal GFP fusion is diffuse when expressed alone but aggregates when co-expressed with CluMPS. Scale bar = 20 µm. Quantification of cluster area **(B)** and fraction of cells with detectable clusters **(C)** of GFP-N in the presence or absence of CluMPS. **(D)** TDP-43 is a nuclear protein whose aggregation is associated with amyotrophic lateral sclerosis (ALS). Scale bar = 20 µm. A GFP-TDP43 fusion is diffuse in the nucleus when expressed alone but forms large clusters when co-expressed with LaG-17 CluMPS fused to an NLS. Quantification of cluster area **(E)** and fraction of cells with detectable clusters **(F)** of GFP-TDP43 in the presence or absence of LaG17-CluMPS(NLS). **(G)** EML4-ALK is an oncoprotein that forms aggregates required for its oncogenic signaling. An EML4-ALK-GFP fusion is clustered in few cells when expressed alone but robustly forms droplets when co-expressed with LaG17-CluMPS. Scale bar = 20 µm. Quantification of cluster area **(H)** and fraction of cells with detectable clusters **(I)** of EML4-ALK-GFP in the presence or absence of LaG17-CluMPS. Box-and-whisker plots in (**B,E,H**) represent median and quartiles from approximately 20-50 cells per group. Vertical lines extend from quartiles ± 1.5*(interquartile range). Bar plots (**C,F,I**) represent frequency. Error bars = 95% confidence interval of approximately 20-50 cells per group

We then tested CluMPS amplification of TDP-43, a nuclear protein that is found in neuronal inclusions in ∼97% of sporadic ALS patients, and whose aggregation is strongly associated with ALS progression^74–76^. Because TDP-43 is a nuclear protein, we appended LaG17-CluMPS with a nuclear localization sequence (NLS) at the C-terminus. GFP-TDP-43 was expressed in the nucleus and, like N-GFP, was largely diffuse in HEK 293T cells under 40X confocal microscopy (**Figure 4D**). By contrast, co-expression with nuclear-targeted LaG17-CluMPS increased the detection of TDP-43 condensates from ∼2% to ∼80% of cells (**Figure 4D-F**), revealing the underlying self-association tendencies of GFP-TDP-43. Phase separation was more evident in the CluMPS channel, likely an artifact of differential expression levels and background fluorescence of the target and reporter (**Figure 4D**).

Finally, we tested the ability of CluMPS to amplify condensates of EML4-ALK, an oncogenic fusion protein commonly found in lung cancers^74,84^. The EML4-ALK-GFP fusion alone was observed to form condensates in only ∼25% of transfected HEK 293T cells and remained diffuse in the remaining 75% of cells (**Figure 4G-I**). However, co-expression with LaG17-CluMPS yielded large condensates in virtually every cell (**Figure 4G-I**). Thus, in the absence of CluMPS, GFP fusions of EML4-ALK form clusters that are too small to see above the diffuse GFP background in most cells, and CluMPS can successfully detect and amplify these small clusters.

### Visualizing clusters of endogenous proteins

Because CluMPS magnifies clusters of its binding partner, it can be applied to detect clusters of endogenous proteins. This could be achieved either by using binding partners for the unmodified target or by tagging the protein of interest with a custom binding epitope at the endogenous locus. We tested the first strategy by designing a CluMPS reporter to amplify condensates of the EML4-ALK oncogene in patient-derived H3122 lung cancer cells (**Figure 5A**). EML4-ALK drives oncogenic signaling by the cytoplasmic aggregation and autophosphorylation of the ALK kinase domain, which recruits adapter proteins including Grb2 and Gab1 to trigger downstream growth signals^75^. For the CluMPS binding domain, we used the proline rich domain (PRD) from Gab1 (amino acids 263-451), which binds Grb2^85^, an essential component of EML4-ALK condensates^77^ (**Figure 5A,B**). Thus Gab1(PRD)-CluMPS reports on Grb2 condensation as a proxy for EML4-ALK condensation. In H3122 cells, we observed large condensates of Gab1(PRD)-CluMPS (**Figure 5C,D**), whereas no condensation was observed in cells expressing a construct with either no amplification domain (Gab1(PRD)-mCh) or no Grb2-binding domain (LaG17-CluMPS) (**Supplementary** Figure 15A,B). EML4-ALK puncta that co-localized with Gab1(PRD)-CluMPS were larger than those that did not (53 ± 2 vs. 11.0 ± 0.1 pixels, roughly 3.2 ± 0.12 vs 0.66 ± 0.01 μm^2^), indicating successful CluMPS amplification of the endogenous condensates (**Figure 5D, Supplementary** Figure 16A-C). CluMPS expression in H3122 cells that harbored a fluorescently-tagged endogenous allele of Grb2 confirmed that Gab1(PRD)-CluMPS condensates colocalized with both Grb2 and EML4-ALK (**Figure 5E)**. Virtually every CluMPS condensate (97 ± 1%, N = 956 condensates) overlapped with ALK immunostaining, demonstrating high specificity (**Figure 5E**) whereas Gab1(PRD)-CluMPS was diffuse in cell lines that lacked EML4-ALK (**Supplementary** Figure 15C). We observed CluMPS condensates in 89 ± 4 % of CluMPS-expressing cells (N = 224 cells), demonstrating high sensitivity.

**Figure 5.**
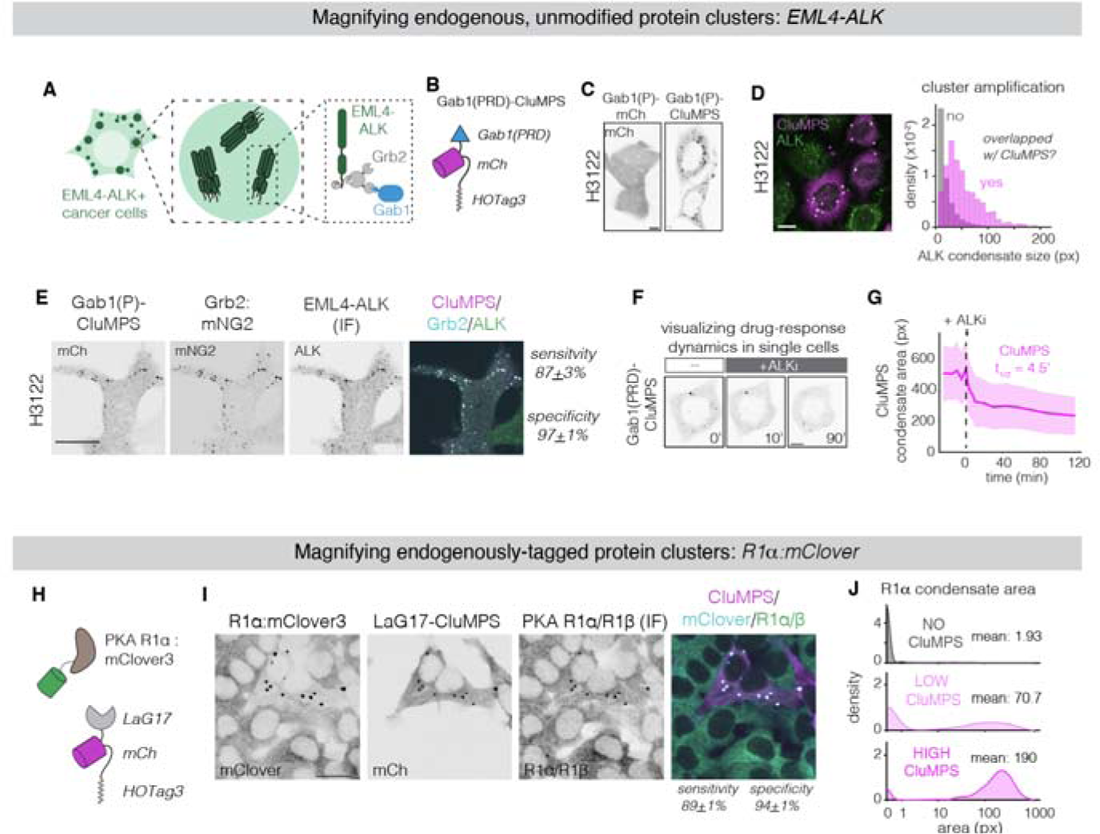
Detection of endogenous clustered proteins using two distinct strategies. **(A)** EML4-ALK forms oncogenic protein condensates that drive oncogenic signaling. Condensation and signaling require the downstream adapter Grb2, which recruits additional adapters including Gab1. **(B)** A CluMPS reporter to visualize endogenous EML4-ALK condensates was developed using the proline rich domain (PRD) from Gab1, which binds an SH3 domain of Grb2 (Gab1(PRD)-CluMPS). **(C)** Gab1(PRD)-CluMPS shows robust condensates in living EML4-ALK+ cancer cells (right). A Gab1(PRD)-mCh fusion is insufficient to visualize EML4-ALK condensates. Scale bar = 10 µm. **(D)** Immunostaining of ALK in H3122. Condensates are larger in cells containing CluMPS (see also **Supplementary** Figure 16). Scale bar = 10 µm. Plot shows distribution of cluster sizes (pixels) for EML4-ALK condensates that are colocalized with a CluMPS condensate (magenta) or not (grey). Data represents approximately 90,000 non-overlapped condensates and 1000 overlapped condensates. **(E)** Immunofluorescence imaging of Gab1(PRD)-CluMPS, Grb2:mNeonGreen2, and EML4-ALK in H3122 cells harboring endogenously tagged Grb2 (Grb2:mNG2). Condensates are enriched for all three species. Scale bar = 20 µm. **(F)** Live cell imaging of CluMPS condensates before and during ALK inhibition with 1 μm crizotinib (see also **Supplementary Movie 4**). Scale bar = 10 µm. **(G)** Quantification of CluMPS dynamics. CluMPS condensate area remains stable for 30 minutes prior to drug treatment, then rapidly decreases upon treatment with ALKi. Half-lives derived from fitting data between 0 and 90 minutes to a single exponential decay. Trace represents mean. Ribbons = 95% CI of 31 cells. **(H)** R1α, a regulatory subunit of the PKA holoenzyme, was tagged at the endogenous locus with mClover. (see **Supplementary** Figure 18). LaG17, which has high affinity for mClover, can be used in a CluMPS reporter (LaG17-mCherry-HOTag3) to report on clusters of PKA R1α via binding the fused mClover tag. **(I)** Images of fixed HEK 293T with mClover-tagged R1α with some cells transfected with CluMPS and stained for R1α/β. Cells with CluMPS have large condensates, while cells lacking CluMPS appear to have diffuse R1α. Condensates are enriched in all three species. LaG17-CluMPS showed high sensitivity against cells with mClover-tagged R1α (89% cells with CluMPS condensate) and specificity (94% of CluMPS condensates are colocalized with an R1α/β condensate). Scale bar = 20 µm. **(J)** Density plots show distribution of condensate area in cells with no CluMPS (N = 2832), low CluMPS (N = 2428), or high CluMPS (N = 743), with text showing mean area of condensates (pixels) in all cells in each group.

The ability to visualize endogenous aggregates allowed us to track their dynamics in response to the ALK inhibitor crizotinib (**Figure 5F, Supplementary** Figure 16D-H**, Supplementary Movie 4**). Although it is known that crizotinib blocks EML4-ALK activity and suppresses its condensation, in part due to dissociation of Grb2^58^, the kinetics and extent of condensate dissolution has not been observed in living cancer cells. We found that, upon drug addition, CluMPS condensate area rapidly decreased within 10 minutes, followed by a slow but measurable decrease thereafter (**Figure 5G, Supplementary** Figure 16D-H). The half-life of the initial decay of the CluMPS signal was ∼4 minutes. This rate is ∼4X faster than a recent measurement of the rate of Grb2 dissociation from heterologously overexpressed EML4-ALK^78^. We validated this fast decay through a fixed-cell time course experiment and obtained similarly fast decay of the EML4-ALK/CluMPS condensates (t_1/2_ = 3.5 minutes) (**Supplementary** Figure 16G,H**, Supplemental Figure 17**). This faster measured rate may reflect the differences in measuring endogenous aggregates versus aggregates of overexpressed EML4-ALK, as performed previously^57^. We also observed that, despite extended drug treatment, ALKi did not induce full dissociation of CluMPS condensates, in line with previous reports showing that EML4-ALK (V1) condensates are relatively stable^61^. These residual CluMPS condensates were not merely an artifact of CluMPS expression, since residual ALK clusters were also detected in untransfected, drug-treated H3122 cells (**Supplementary** Figure 16B-D). The presence of CluMPS condensates under these drug conditions suggests that the residual EML4-ALK clusters retain a basal level of phosphorylation and signaling potential, which is required for the colocalization of Grb2 and Gab1(PRD). Thus, CluMPS can visualize the dynamics of endogenous protein assemblies and showcases potentially important differences between measuring endogenous proteins in their naive context as compared to their exogenous overexpression in unrelated cell types.

We next tested whether CluMPS could also amplify clusters of proteins that have been tagged at their endogenous locus. Phase separation of the protein kinase A (PKA) regulatory subunit R1α was recently found to sequester intracellular cAMP and regulate PKA signaling ^20^. When overexpressed, R1α forms large condensates, but endogenous clustering is more difficult to visualize ^20^. To apply CluMPS to endogenous R1α, we first tagged R1α with mClover3 at its endogenous locus in HEK 293T cells (**Figure 5J**, **Supplemental Figure 18**) ^75^. Because mClover3 is a GFP derivative, it served as an affinity tag for LaG17-CluMPS (**Figure 5H**).

Whereas R1α:mClover3 looked diffuse under 40X magnification, co-expression of LaG17-CluMPS yielded large condensates containing both CluMPS and mClover3 (**Figure 5I,J**). Immunofluorescence confirmed that these condensates were enriched for R1α/β (**Figure 5I**). As before, CluMPS detection was sensitive (condensates were present in 89% of CluMPS positive cells, N = 743 cells), and specific (94% of CluMPS condensates were enriched for R1α/β, N = 1302 condensates). Endogenous tagging presents a straightforward strategy for amplifying clusters of endogenous proteins, providing a generalizable strategy for using CluMPS to report on arbitrary endogenous proteins with no further engineering of the reporter.

### Multiplexed amplification of distinct target clusters in single cells

Finally, we asked whether we could multiplex CluMPS reporters to magnify distinct clusters in the same cell. We leveraged the fact that both HOTag3 and HOTag6 work as strong amplification domains (**Figure 1, Supplementary** Figure 1) but do not cross-interact when co-expressed ^63^.

We generated a HOTag6-based CluMPS reporter using miRFP and LaG17 as the binding domain (LaG17-CluMPS) to multiplex with a HOTag3-based reporter with mRuby2 and a nanobody for the ALFA tag^75^ (nbALFA-CluMPS)(**Figure 6A**). We co-expressed these two constructs in cells with two distinct optogenetic clustering targets: ALFA-Cry2, which forms clusters in the cytoplasm, or BcLOV4-GFP, which forms clusters at the membrane (**Figure 6B**)^77^. Blue light triggered the appearance of distinct condensates in both the mRuby and iRFP channels, reflecting amplification of the two distinct optogenetic clusters (**Figure 6C,D, Supplementary Movie 5**). Condensates in the two channels did not overlap, with membrane-associated BcLOV4 clusters more peripheral and Cry2 clusters more cytoplasmic, thus demonstrating multiplexed cluster detection using orthogonal CluMPS probes (**Figure 6C,E, Supplementary** Figure 19).

**Figure 6.**
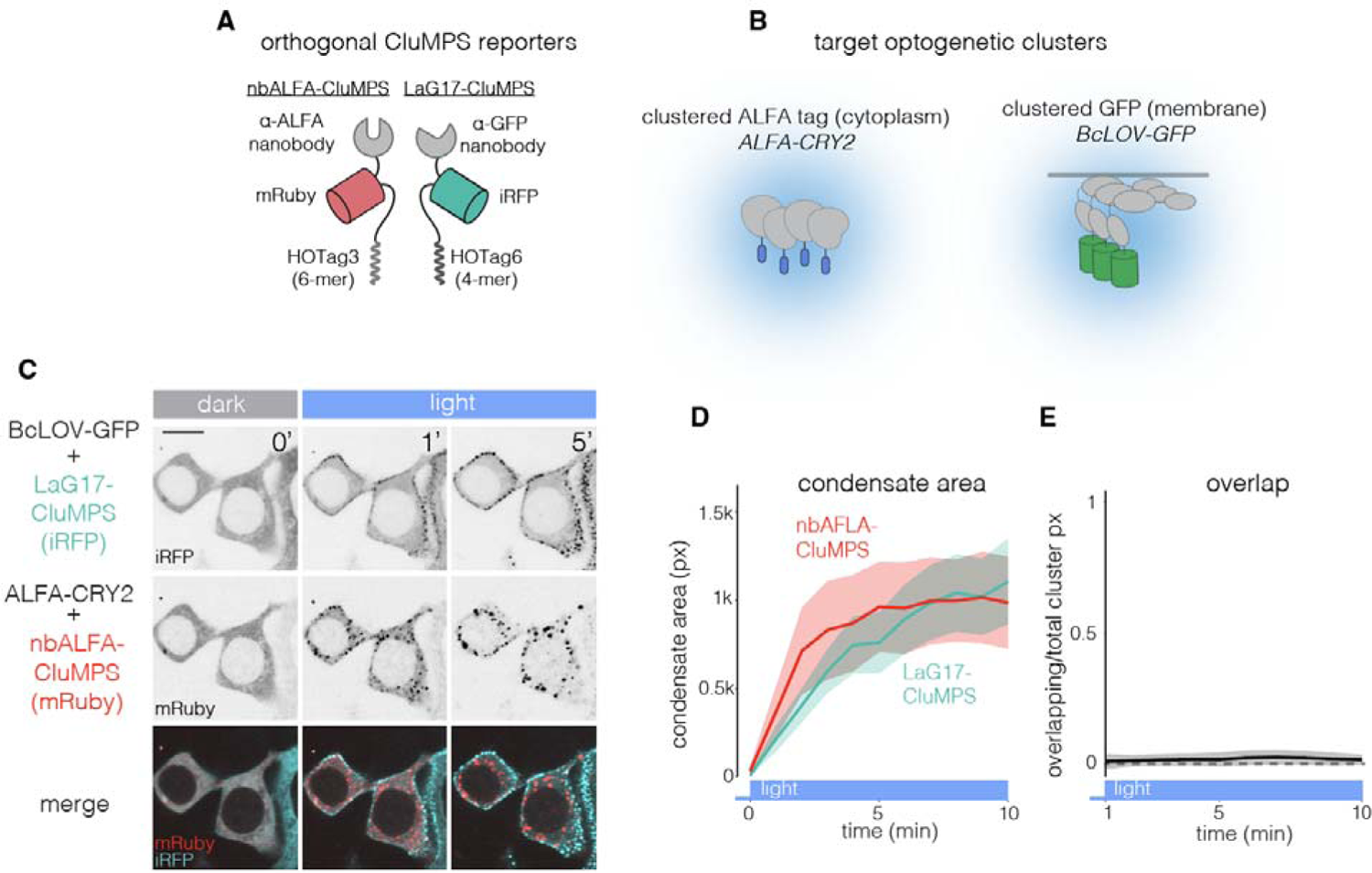
Multiplexed imaging of orthogonal CluMPS reporters. **(A)** Orthogonal CluMPS reporters can be generated to visualize multiple targets in the same cell. **(B)** Cry2 and BcLOV4 are two photosensitive proteins. Cry2-GFP forms multimers in the cytoplasm, while BcLOV4 clusters and translocates to the plasma membrane. **(C)** In cells expressing both targets (ALFA-Cry2 and BcLOV4-GFP) and reporters (nbALFA-mRuby-CluMPS and LaG17-iRFP-CluMPS), both reporters are diffuse in dark-adapted cells. Blue light stimulation of target clustering triggers non-overlapping CluMPS condensation in both channels. Scale bar = 20 µm. **(D)** Quantification of area of nbALFA-CluMPS (red) and LaG17-CluMPS (teal) clusters per cell over 10 minutes of blue light stimulation. Data represent mean, ribbons = 95% CI of 25 cells that expressed all components. **(E)** Quantification of overlap between the two CluMPS channels during light stimulation. Trace represents mean, shading represents 95% CI of 25 cells.

## Discussion

The CluMPS reporter visualizes small protein clusters by amplifying them, with a sensitivity that extends to small oligomers (3-4-mers) under appropriate conditions. Through simulations and experiments, we showed that amplification is a function of concentration, affinity for the target, target cluster size, and the fraction of target that is clustered. Although our work focused on amplification of small clusters, CluMPS works over a large range of cluster sizes and thus will find use to visualize a wide variety of protein assemblies. We demonstrate that CluMPS remains diffuse when binding monomeric targets, allowing users to be confident that condensation of CluMPS happens only when the CluMPS target is genuinely multimeric. Our CluMPS reporters that bind GFP or ALFA-tag can be used to report on fusions of these tags —both heterologous and endogenous — with arbitrary proteins of interest. Alternatively, CluMPS can be generated to visualize the cluster dynamics of unmodified endogenous proteins and, with an activity-gated binding domain, even downstream signaling (as with Gab1(PRD)-CluMPS, **Figure 5**). Finally, we showed that the CluMPS reporter can be multiplexed using orthogonal CluMPS variants to visualize multiple clusters in individual cells simultaneously.

CluMPS provides several advantages relative to existing methods to visualize aggregates in cells. Unlike FRET, fluorescence anisotropy, fluorescence correlation, or super-resolution techniques, CluMPS can be imaged using a single channel, has high signal-to-noise ratio, and can be observed without complex microscopy and image processing. Further, CluMPS does not require overexpression of the target protein and thus can avoid altering target concentration levels, providing confidence that the detected clusters were in fact present in their physiological context. A recent study developed a transcriptional reporter of protein aggregation called yTRAP, where the aggregation of a protein of interest sequestered a transcriptional activator and reduced fluorescence from a transcriptional cassette^86^. However, this approach required large, stable aggregates of overexpressed protein and could only report on aggregates over long timescales (∼hours to days) in yeast. By contrast, CluMPS allows detection of endogenous, dynamic protein clustering across a broad range of cluster types and sizes within the native cellular context of mammalian cells (**Figure 3**).

A main limitation of CluMPS is that a negative result (no condensates) does not necessarily indicate a lack of target clusters, but could alternatively be explained by factors including low binding affinity for the target, low fraction of clustered target, or mismatch between reporter and target expression. Because some of these factors will vary from cell to cell, CluMPS also cannot be used for ‘back-calculation’ of features like the precise valency of the underlying target cluster. Thus CluMPS is best interpreted as a ∼binary cell-level reporter of a protein’s cluster status.

Nevertheless, our simulations and experiments set forth design rules for CluMPS reporters and provide context for interpreting CluMPS activity, for example when targeting endogenous proteins whose cluster status and distribution are unknown. As is good practice for any live cell reporter, conclusions from CluMPS should be validated by an orthogonal approach (e.g. immunofluorescence). Another limitation is that CluMPS necessarily perturbs the clustering state of its targets, changing the size and potentially the dynamics or even interaction partners of the target clusters. Thus, proper controls are essential to account for these possibilities.

Nevertheless, we have not observed ‘false positive’ CluMPS signals where CluMPS would induce target clustering where none existed, nor even substantial changes in the rates of cluster formation or dissolution (**Supplementary** Figures 2,3**,5,6,9,15**). Finally, although we used CluMPS probes that incorporate mCherry, mRuby, GFP, and miRFP, the identity of the fluorescent protein can have a strong effect on clustering and phase separation of the construct^36,87^. Thus, the suitability of CluMPS variants with different fluorescent proteins must be determined empirically.

We anticipate that the simplicity and high signal-to-noise ratio of CluMPS will enable numerous unexplored applications, for example for screening drug libraries for compounds that disrupt small neurodegenerative aggregates, or for detecting pathogenic aggregates in patient biopsies. We thus expect CluMPS will enhance our understanding of the prevalence, relevance, and dynamics of protein clusters that have previously been hard to see.

## Acknowledgements

We thank Alex Hughes (Penn), Matthew Good (Penn) and Jin Zhang (UCSD) for helpful discussions. We also thank Matthew Good and Ophir Shalem for providing genetic constructs used in this work. This work was supported by the National Institutes of Health (R35GM138211 for L.J.B and D.G.M.; R35GM146877 for J.G.). Cell sorting was performed on a BD FACS Aria Fusion that was obtained through NIH S10 1S10OD026986.

## Author Contributions

T.R.M. and L.J.B conceived the study; T.R.M., A.I., D.G.M, A.A.P designed and performed experiments; D.R. built the kinetic Monte-Carlo model; T.R.M. wrote the script for automated analysis of fluorescent puncta; E.R. conceived and supervised the FCS experiments, which were performed and analyzed by E.B; M.C.C. and J.G. generated the Grb2:mNG H3122 cell line. T.R.M. analyzed data; T.R.M. and L.J.B. wrote the manuscript and made figures; L.J.B. supervised the work.

## STAR Methods

### RESOURCE AVAILABILITY

#### Lead Contact

Requests can be made to the lead contact Lukasz Bugaj.

#### Materials Availability

Select plasmids from this manuscript have been deposited on addgene (https://www.addgene.org/Lukasz_Bugaj, ID 190024, ID 190026)

#### Data and Code Availability

● All microscopy data available from the lead contact upon request.
● Code used to fit clusters throughout this manuscript and described in Supplementary Figure 20 and 21 is available on github (https://github.com/BugajLab/Cluster-Fitting, DOI: 10.5281/zenodo.10471925).
● Any additional information required to reanalyze the data reported in this paper is available from the lead contact upon request.

## EXPERIMENTAL MODEL AND STUDY PARTICIPANT DETAILS

### Cell lines and cell culture

All cell lines were maintained in standard cell culture incubators at 37°C and 5% CO_2_. Lenti X HEK 293T cells and HeLa cells were cultured in DMEM containing 10% fetal bovine serum (FBS) and 1% penicillin/streptomycin (P/S). H3122 cells were cultured in RPMI-1640 media containing L-glutamine with 10% FBS and 1% P/S. For experiments, cells were seeded with P/S-free media in 96 or 384 well plates coated with 10 µg/mL MilliporeSigma™ 597 Chemicon™ Human Plasma Fibronectin Purified Protein in PBS. For 384 well plate experiments, 3000 HEK 293T cells or H3122 cells were seeded in 50 μL of cell culture media in each well. For 96 well plate experiments, 20,000 HEK 293T or H3122 cells were seeded in 200 μL of cell culture media in each well.

## METHOD DETAILS

### Plasmid design and assembly

Constructs were assembled using Gibson assembly. DNA fragments and backbones were generated via PCR and inserted into the backbone via HiFi cloning mix (New England Biolabs). DNA sequences encoding GFP binding nanobodies (LaGs) were a kind gift from Dr. Michael P. Rout ^79^. HOTag6 and HOTag3 were obtained from pcDNA3-ERK-SPARK, which was a gift from Xiaokun Shu (Addgene plasmid # 106921; http://n2t.net/addgene:106921; RRID:Addgene_106921) ^80^. FUS(LC) (1-163), Xvelo, and LAF-1 RGG were kindly provided by Matthew C. Good. iLid strings were adapted from mCherry-iLidx6, which was a gift from Takanari Inoue (Addgene plasmid # 103779; http://n2t.net/addgene:103779; RRID:Addgene_103779) ^81^. GFP-TDP-43 was kindly provided by Ophir Shalem. SARS-CoV2-N protein sequence was sourced from pLVX-EF1alpha-SARS-CoV-2-N-2xStrep-IRES-Puro, which was a kind gift from Nevan Krogan (Addgene plasmid # 141391; http://n2t.net/addgene:141391; RRID:Addgene_141391). EML4-ALK-GFP was made through fusing EML4-ALK(V1) to GFP as previously described ^82^. FTH1 was sourced from pCMV-SPORT6_FTH1, obtained from the High-Throughput Screening (HTS) Core at Penn Medicine. The sequence encoding ALFA tag was synthesized as a single strand DNA oligomer (Genewiz from Azenta Life Sciences), and sequence encoding the ALFA binding nanobody (peptide elution, “nbALFA(PE)” variant) was synthesized as a gBlock gene fragment (Integrated DNA Technologies) ^83^. Gab1 (PRD) was sourced from the human GAB1 cDNA sequence obtained from the High-Throughput Screening (HTS) Core at Penn Medicine. PCSDest vectors for RNA synthesis were obtained from the Zebrafish Core at Children’s Hospital of Philadelphia. miRFP670, used in iLid_strings_ and LaG17-miRFP670-HOTag6, was adapted from pCSII-EF-miRFP670v1-hGem(1/110), which was a gift from Vladislav Verkhusha (Addgene plasmid # 80006; http://n2t.net/addgene:80006; RRID:Addgene_80006).

### RNA synthesis

RNA transfection was used instead of DNA transfection for cell types that showed low efficiency of plasmid DNA transfection. To generate RNA from the PCSDest expression plasmid, the plasmid DNA was linearized with NotI and used as a template for RNA synthesis. RNA was synthesized using the mMESSAGE mMACHINE™ (ThermoFisher Scientific) transcription kit with SP6 RNA polymerase. RNA was spin-column purified using RNA Clean and Concentrator kit (Zymo Research).

### Plasmid and RNA transfection

HEK 293 T cells were transfected with DNA plasmids using Lipofectamine™ 3000 Transfection Reagent (ThermoFisher Scientific) per manufacturer’s protocol. Final transfection mixture was 1-30 ng μL^-1^ DNA, 2% Lipofectamine™ reagent, 2% P3000 reagent and was brought to final volume with Opti-MEM™ (ThermoFisher Scientific). Cells seeded in 384 well plates received between 1 and 5 μL of transfection mixture, and cells in 96 well plates received 10 μL of transfection mixture. Cells were imaged 24-60 hr after transfection. H3122 cells were transfected with RNA using Lipofectamine™ MessengerMAX™ Transfection Reagent (ThermoFisher Scientific), as RNA transfection was found to be more efficient than DNA transfection (described above) in this cell type. Final transfection mixture was 10 ng μL^-1^ RNA, 2% Lipofectamine™ MessengerMAX™ reagent, and was brought to final volume with Opti-MEM™. Cells were imaged 12-36 hr after RNA transfection. HEK 293 T cells and HeLa cells were also transfected with Gab1(PRD) CluMPS RNA (as described above) for comparison with H3122 cells in **Figure S15**.

### Live cell imaging

Live-cell imaging was done using a Nikon Ti2E microscope equipped with a Yokagawa CSU-W1 spinning disk, 405/488/561/640nm laser lines, an sCMOS camera (Photometrics), a motorized stage and an environmental chamber (Okolabs). HEK 293 T cells were imaged with a 40X oil immersion objective at 37°C and 5% CO_2_. In addition to exposure for imaging, experiments requiring optogenetic stimulation of Cry2 and/or iLid-sspB received an additional 500 ms per field of view per minute blue light exposure using the 488 nm laser line. For experiments where HEK 293 T cells were transfected with a construct containing miRFP670, the media was supplemented with 25 μM biliverdin hydrochloride (Sigma-Aldrich, product number 30891) 24 hours prior to imaging. H3122 cells were imaged with a 40X oil immersion objective. H3122 cells were starved overnight with RPMI-1640 media containing L-glutamine and 1% P/S using a BioTEK 405 plate washer. For live nuclear staining, H3122 cells were treated with 5 µg/ml Hoechst 33342 for 15 min before imaging. Cells were imaged for 30 minutes, then treated with 1µM crizotinib (Sigma-Aldrich, PZ0191) and imaged for 2 hours.

### Immunofluorescence staining

Immediately after treatment, cells were fixed and stained as previously described ^88^. Briefly, cells were fixed for 10 min in 4% paraformaldehyde. Cells were permeabilized using PBS with 0.5% Triton-X100 for 10 min. Cells were further permeabilized using 100% methanol at −20°C for 10 min and samples were blocked with 1% bovine serum albumin (Fisher, BP9706100) in PBS + 1% BSA for 1 hour at room temperature. Cells were then incubated in primary antibody diluted in PBS + 1% BSA for 2 hours at room temperature or overnight at 4 °C (anti-ALK (D5F3), Cell Signaling, catalog number 3633, 1:400 for ALK staining and anti-R1α/β, Cell Signaling, catalog number 3927, 1:100 for R1α/β staining). Primary antibody was then washed with 0.1% Tween-20 in PBS (PBS-T). Cells were then incubated in secondary antibody diluted in PBS + 1%BSA as recommended (IgG (H+L) Cross-Adsorbed Goat anti-Rabbit, DyLight™ 488, Invitrogen, catalog number 35553, 1:500) and 4,6-diamidino-2-phenylindole (DAPI; ThermoFisher Scientific, catalog number D1306, 300 nM) for 1 hour at room temperature and washed with PBS-T. Cells were imaged in 50 µL of PBS.

### iLid strings experimental procedures

For testing CluMPS sensitivity with small multimers using iLid_strings_ (multiple iLid domains fused on a single peptide, iLidxN: N = 2:6 iLids fused to miRFP670) transfected with GFP-sspB, it was important to achieve a broad range of transfection conditions in order to sample a range of sspB:iLid ratios across cells (Figure 3F-H). For these transfections, plasmids were ‘poly-transfected’; each plasmid was mixed with transfection reagents in a separate tube from the other plasmids and added to cells separately (**Supplementary** Figure 11A)^70^. This resulted in uncorrelated expression between components, permitting sampling of a wide range of sspB:iLid ratio across cells (**Supplementary** Figure 11B).

### iLid_string_ analysis

To determine the stoichiometric ratio of sspB:iLid, which sets the fraction of EGFP-sspB that could bind sites on iLid multimers vs the fraction that remains monomeric, we used fusion of EGFP-miRFP670 as a control (**Supplementary** Figure 10A). For each iLid_string_ experiment, a separate well on the same plate was transfected with this EGFP-miRFP670 control, and imaged with the same microscope parameters as the wells containing EGFP-sspB, miRFP670-iLid_string_, and LaG17-CluMPS. With this control, the ratio EGFP:miRFP670 intensity in cells transfected with EGFP-miRFP670, in which the number of EGFP and miRFP670 proteins are assumed to be equal, could be calculated for many cells (**Supplementary** Figure 10B,C). The median ratio is then used as a normalization factor to determine the stoichiometric ratio of EGFP:miRFP670 in cells transfected with EGFP-sspB and miRFP670-iLid_string_ (**Supplementary** Figure 10C,D). With this, it is trivial to calculate the sspB:iLid ratio given that EGFP-sspB are 1:1 and miRFP670:iLid is known and dependent on the valency of the specific iLid_string_ that was transfected. The percentage of GFP-sspB that was able to participate in clustering can then be estimated as min(100, 100*(iLid/sspB)).

### Estimating true concentration of nbALFA-GFP-HOTag3 and iRFP-ALFAx4

Imaging of 1)purified GFP of known concentration and 2) cells transfected with GFP-iRFP was used to estimate the concentration of nbALFA-GFP-HOTag3 and iRFP-ALFAx4 in cells transfected with both constructs (**Supplementary** Figure 14B). Wells not containing cells (but on the same plate as the cells) were filled with 40 μL of purified GFP (abcam, AB84191) at concentrations of 25 μM, 10 μM, 5 μM, 2.5 μM, 1 μM, and 0.5 μM and imaged to obtain the relationship between GFP intensity and GFP concentration (**Supplementary** Figure 14C). Also on the same day and plate, cells transfected with GFP-iRFP were imaged to understand GFP and iRFP relative intensities when present in equimolar amounts (See previous methods paragraph, **Supplementary** Figure 10). With these two conversion factors (GFP intensity:GFP concentration and GFP intensity:iRFP intensity), the concentration of GFP and iRFP in cells can be estimated from their respective intensities.

### Tagging R1α at the endogenous locus

PKA regulatory subunit 1α (PRKAR1A) was tagged at the endogenous locus using homology independent intron tagging described by Serenrenik et al ^81^. HEK293T cells were seeded in 12 well plates at a density of 250,000 cells per well. Each well was transfected 24 hours after seeding with 467 ng of mClover3 generic donor plasmid, 140 ng of plasmid encoding the sgRNA which targets the donor, 140 ng of plasmid encoding the sgRNA targeting PRKAR1A intronic sequences, 252 ng of plasmid encoding Cas9. These plasmids were a kind gift from Dr. Ophir Shalem, and were unmodified save for the insertion of the sgRNA sequence targeting PRKAR1A. sgRNA sequences available in Key Resource Table. 24 hours after transfection, cells were passed to a 6 well plate and seeded in media containing 5 μg/mL blasticidin. After two weeks of blasticidin selection, cells were sorted (BD FACS Aria Fusion) to further enrich fluorescent-positive cells. Single cell colonies were made by seeding cells on 96 well plates with a density of 0.5 cells/well to ensure wells contained 0-1 cells. Colonies were screened via staining for PKA R1α/R1β (cell signaling #3927) and validated by western blot (**Supplementary** Figure 18).

### Western blot validation of PKA R1α:mClover tagging

Wild-type and R1α:mClover HEK293T cells were grown to confluency in a 10cm plate. Cells were trypsinized, counted, and washed in room temperature PBS. 4×10^6^ of each cell type were lysed in 500 μL of lysis buffer (50mM Hepes 7.5, 10% glycerol, 150mM NaCl, 1% Triton X-100, 1mM EDTA, in H_2_O). Lysis buffer was supplemented with a protease inhibitor just prior to lysis. 20 μL of lysis was subjected to SDS–polyacrylamide gel electrophoresis (SDS-PAGE). Protein separations were transferred onto a nitrocellulose membrane using the Trans-blot Turbo RTA transfer kit (Bio-rad, #170-4270) according to the manufacturer’s protocol. Membranes were blocked in 5% milk in Tris buffer saline with 1% Tween-20 (TBS-T) for 1 hour and incubated overnight with gentle shaking at 4°C with primary antibody against R1α (CST #5675). Primary antibody was used at a dilution of 1:1000 in TBS-T with 5% BSA. After washing with TBS-T, membranes were incubated with secondary antibodies in TBS-T with 5% milk for 1 hr at room temperature (Alexa Fluor® 790 AffiniPure Donkey Anti-Rabbit IgG (H+L) Jackson # 711-655-152). Membranes were then imaged on the LI-COR Odyssey scanner.

### Fluorescence correlation spectroscopy

For in-cell fluorescence correlation spectroscopy (FCS) measurements, HEK 293T cells were plated in fibronectin coated 8-well NUNC chambers (Thermo Scientific, Rochester, NY) at a cell density of 60,000 cells per well. Cells were transfected with 50 ng of GFP or GFPx5 and then exchanged into 250-µL phenol red-free media 24-hr after transfection. A portable stage-top incubator (37 °C and 5% CO_2_) was used during measurements. Imaging and FCS measurements were made 24-72 hours after transfection.

Fluorescence lifetime imaging (FLIM) and FCS measurements were carried out on a PicoQuant MicroTime 200 fluorescence microscope system equipped with a Flimbee Galvo Scanner to allow for imaging (PicoQuant, Berlin, Germany). A 482 nm excitation laser set to 65% of its maximum output, a pulse rate of 40 mGHz, was focused by a 60x Plan-Apo/1.4-NA water-immersion objective (Olympus, Tokyo, Japan) with the coverslip correction collar set to 0.15.

Fluorescence emission was collected through the objective and focused onto a 150 μm diameter pinhole, then directed through a 525 ± 35 nm band-pass filter and finally collected by an avalanche photodiode detector. The Nunc chambers were mounted to a scanning stage composed of a manually adjustable XY-axis stage, a Nano-ZL 100 piezo scanning stage (Mad City Labs, Madison, WI). The Z-piezo stage and galvo-scanner allow for precision 2-dimensional imaging of the X/Y, X/Z, and Y/Z planes. Image acquisition, data collection, and autocorrelation analysis of intensity fluctuations were done using PicoQuant’s accompanying software SymPhoTime 64.

For FCS measurements, cells were first imaged in the X/Y plane and evaluated based on the average photon count rate of the cell; cells with count rates of 150–600 kHz under the conditions of our measurements were selected to move forward. For a selected cell, a better quality FLIM image was first obtained in the X/Y plane, followed by an X/Z or Y/Z image. This image was used to manually select placement of the focused laser in the cell cytoplasm, avoiding positioning too close to the plasma membrane or nucleus. Generally, 3 to 5 FCS measurements were made for each cell. For each FCS measurement, five autocorrelation curves of five seconds each were collected and averaged together. Using a lab-written script in MATLAB, the averaged autocorrelation curves were fit to a function for a single fluorescent species undergoing Brownian motion in a three-dimensional Gaussian volume, with fits weighted by the inverse square of the standard deviation:

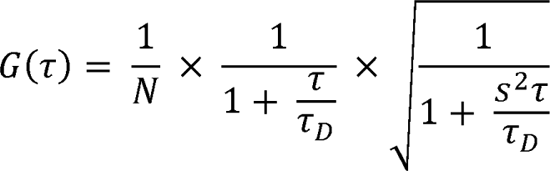

where *N* is the average number of molecules in the focal volume, *s* is the ratio of the radial and D is the translational diffusion time. For each FCS measurement, the molecular brightness of the fluorescent species interrogated – expressed as counts per molecule (CPM) – was calculated by dividing the average intensity by *N*.

### Cell segmentation

Cells were segmented using either CellProfiler for automated segmentation or by hand with a custom MATLAB script. For CellProfiler segmentation, nuclei were identified. Then, cytoplasmic fluorescence channels were gaussian and median filtered to prevent artifacts from segmenting into clustered cells, and nuclear masks were expanded using the watershed method to segment cell cytoplasms. For custom segmentation in MATLAB, a script was developed that allowed a user to manually draw and modify cell masks. This script was used in cases where highly accurate, low-throughput cell segmentation was needed. For quantifying clusters, segmentation masks were fed forward into the cluster fitting algorithm (**see below**).

### Fitting and quantifying clusters

Clusters were quantified from images of live and fixed cells using a custom MATLAB script. After cell segmentation, clusters were quantified in each cell at each timepoint (**Supplementary** Figure 21). The algorithm processes a single cell at a time, first normalizing the intensities within the cell by subtracting the minimum intensity within the cell, then dividing by the median background-subtracted intensity within the cell. Pixel intensities at the edge of the cell mask are then propagated to the surrounding pixels outside of the cell mask to reduce contrast at the edges of the cell (**Supplementary** Figure 20C)).

Selecting which pixels within the cell are included in a cluster is a two step process. First, single high-intensity pixels at the center points of clusters are identified. To select these pixels, a series of transforms is applied to the normalized cell image to remove background, suppress noise, and enhance medium-high frequency objects (clusters). These are as follows:

- Gentle gaussian low-pass filter to suppress the highest frequency noise. This aims to prevent single pixels that are randomly of high intensity from being enhanced by the subsequent contrast enhancing steps.
- A top hat filter to suppress background; areas with low and uniform (longer than length scales of ∼5 pixels) values.
- Another gentle gaussian low-pass filter is used again to suppress the highest frequency contrast, which tends to be noise rather than clusters
- Convolution with a 5×5 grid with the following values

- This is a laplacian edge detector, and is typically used to enhance edges/areas of contrast
- Resultant negative values are set to zero
- Values less than zero occur when a pixel has low intensity compared to its surroundings. Suppression of such features enhances cluster-finding.

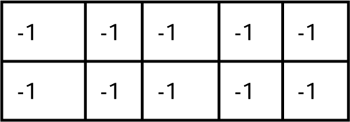

|  |  |  |  |  |
| --- | --- | --- | --- | --- |
| -1 | -1 | 24 | -1 | -1 |
| -1 | -1 | -1 | -1 | -1 |
| -1 | -1 | -1 | -1 | -1 |

With this enhanced contrast image (**Supplementary** Figure 20E,G), points above a user-defined threshold are identified. Quality control is performed to delete pixels from this list until only the highest intensity pixel (in the raw fluorescent image, rather than the transformed image) from each cluster is kept. This is defined as the cluster center (**Supplementary** Figure 21C).

Next, an expansion algorithm determines which pixels are included in the cluster. From each identified cluster center, adjacent pixels are checked to be included or excluded from the cluster as determined by an intensity cutoff. The cutoff is set by the local background in the area surrounding the cluster or by the median of the cell (whichever is lower) multiplied by a scaling factor set by the user. Pixels above this cutoff are included in the cluster, while pixels below the cutoff are discarded (**Supplementary** Figure 21D). This occurs recursively, where adjacent pixels to each included pixel are also checked to be included or excluded until all new neighbors are below the threshold.

With this two step algorithm, we were able to extract information from cells on the single cluster level, where the location, size, and intensity of each cluster in a cell is known.

### Quality control in analysis of cluster fitting from automated fits

When clusters were fit on cells that were segmented in CellProfiler, rare errors occurred where either a large fraction of a cell mask would be background or a mask would contain a cell, and also a portion of a nearby and far brighter cell. In both cases, the algorithm would fit the entire bright portion (either cell in a mask of mostly background, or the portion of the very bright cell segmented with a dimmer cell) as a cluster. To address this, clusters fit from CellProfiler-generated masks were discarded if they were greater than 500 pixels in area, as such clusters were nearly exclusively caused by erroneous fits.

### Analysis of Cry2-GFP + CluMPS variants

In all experiments where Cry2-GFP is expressed alone or with a CluMPS variant, cells are gated for low expression of Cry2-GFP that do not form visible clusters after blue light stimulation. This is demonstrated in Figure 1.

### Data visualization

Clustering data were exported from MATLAB. Data were organized and visualized in Rstudio using the tidyR package ^84^.

### Curve fitting

Gab1(PRD)-CluMPS cluster area quantification during treatment with crizotinib (Figure 5**, Supplementary** Figure 16) were fit to exponential decay functions of the form

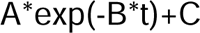

where t is time in minutes, and with the moment of crizotinib addition being t = 0 minutes. A MATLAB function was written where the output was the sum of the squared difference between an exponential decay of the form above and experimental data. Minimization of this function generates the parameters for an exponential decay which most closely match the experimental data. This process required ‘guess’ parameters to act as a starting point for the minimization process. Guess parameters were set as A = the first experimental datapoint minus the lowest experimental data point, B = log(2)/3.5, and C = the lowest experimental data point. Identical methods were used to fit decay of CluMPS condensate area and number in **Supplemental** Figure 6.

### Modeling

We generated a stochastic model of CluMPS amplification of target clustering in Python, using the NumPy^85^, Math, Statistics^86^, and matplotlib^87^ libraries. Our model modifies a previously reported model by Dine et al that describes clustering of a single protein species^24^. We extended this model to simulate the kinetics of condensation of two multivalent species, the key principle that drives CluMPS reporter activation upon clustering of a protein target, using a rejection kinetic Monte Carlo process. The simulation contains two species, the target and reporter. In each simulation there are 350 reporter units and 350 target units. At the simulation’s initialization, targets and reporters are randomly arranged on a 70 x 70 square grid. Each square on the grid can contain up to one target and one reporter, but cannot contain two units of the same type.

Thus, the simulation can be conceptualized as two superimposed planes: one containing targets and one containing reporters (Figure 2A). The model simulates kinetics through accepting or rejecting movements of units on the grids (a random unit of either type and direction for it to move are chosen at each step of the simulation). Whether a movement trajectory is accepted or rejected is determined by a calculated probability for the move to happen, and a random number roll (e.g. a move with a calculated probability of 60% is accepted if a randomly generated number between 0 and 1 is less than 0.6, while the move is rejected if the number generated is greater than 0.6). The probability of a move being accepted is calculated as a function of the enthalpy of the interactions a unit experiences at the initial state when a move is proposed. Units can only interact with units of the opposite species that are at the same or adjacent coordinates (Figure 2A**,B**). Both reporter and target units have a user-defined valency, which determines the maximum number of cross-species interactions a unit is able to participate in. The reporter has a valency of 6 in all simulations, while the target valency varies between simulations (between 2-6) as indicated (Figure 2C**,D****,E,F**). The valency of all units stays constant throughout the duration of a simulation. Interactions between units have a user-defined interaction energy, ΔE. The probability for a trajectory that moves a protein to a position not occupied by another of its type is where T is the sum of the interaction energies from all the interactions that the protein currently experiences (the minimum of the number of interactions and the units valency, multiplied by the energy of a single interaction), following metrics are calculated every 2000 proposed steps: the total interaction strength of the system (the sum of the enthalpies that each protein is currently experiencing), the percent clustering (percent of reporter units that are in a cluster of six or more reporters), and finally the rate of change of the total interaction strength from the previous interval to the current one. After 200 intervals have passed (200 intervals*200 steps per interval = 40,000 steps) the model also sums the last 200 rates of change over their corresponding total interaction strengths. When this sum is <= 0.01, the simulation has reached steady state and ends, and the average percent clustering over the last 100 intervals is reported.

For simulations in Figure 2F, in addition to multivalent target units, additional monovalent target units were added to the simulation. For example, to run a simulation with 50% of clustered target, 350 monovalent target units would be added to the simulation in addition to the 350 multivalent target units. Monovalent targets have the same interaction strength, ΔE, with reporter units, can be randomly selected to move like any other unit, but can interact with a maximum of one reporter unit. When a monovalent target is selected to move to a grid square containing a multivalent target or vice versa, the probability that the two units switch places on the grid is set as *e(-T_1_+T_2_)* interact,ions for the first unit, where *T_1_* is the enthalpy of interactions for the first unit, *T = min(valency, # of neighbors) * E* and *T*_2_ is the enthalpy of interactions for the second unit, calculated in the same way.

### Supplementary Movie Captions

**Supplementary Movie 1. Screening amplification domains for CluMPS reporter.** Representative time lapse videos of candidate CluMPS variants (magenta) co-transfected with Cry2-GFP (green) and stimulated with blue light. CluMPS variants harbor different amplification domains as indicated. Time is mm:ss. Scale bar = 20μm.

**Supplementary Movie 2. Representative simulations of stochastic model of CluMPS amplification.** Each row shows motion of the target and reporter units under the indicated simulation parameters. All rows have identical simulation parameters except for the valency of the target, which is indicated to the left of each row. The right-most column shows the extent of reporter clustering and the total interaction strength of the simulation. Simulations were stopped once the interaction strength reached steady state. See **Methods** for more model details.

**Supplementary Movie 3. CluMPS:target affinity regulates CluMPS amplification.** LaG-CluMPS variants with differing affinities were each co-transfected with Cry2-GFP and stimulated with blue light. Higher-affinity CluMPS provided greater cluster amplification. Time is mm:ss. Scale bar = 20μm.

**Supplementary Movie 4. Gab1(PRD)-CluMPS visualizes endogenous EML4-ALK condensates and their shrinkage in response to ALK inhibitor.** Gab1(PRD)-CluMPS was expressed in H3122 cancer cells, which harbor the EML4-ALK (V1) oncogene. ALK inhibitor crizotinib (1 µM) is added at t = 0 min. Scale bar = 20μm.

**Supplementary Movie 5. Multiplexed imaging of orthogonal CluMPS reporters against distinct clustering targets in single cells.** Orthogonal CluMPS variants (LaG17-CluMPS, nbALFA-CluMPS) were co-expressed with orthogonal blue-light-induced clustering systems (BcLOV4-GFP and ALFA-Cry2) in the single cells. Blue light is applied at the beginning of the video to induce clustering of both GFP and ALFA-tag. Distinct clusters are observed in both the miRFP (LaG17 CluMPS, teal) and mRuby2 (nbALFA CluMPS, red) channels, suggesting independent amplification of the distinct clusters. Time is mm:ss. Scale bar = 20μm.

