## Supplementary Figures for "Simple visualization of submicroscopic protein clusters with a phase-separation-based fluorescent reporter"

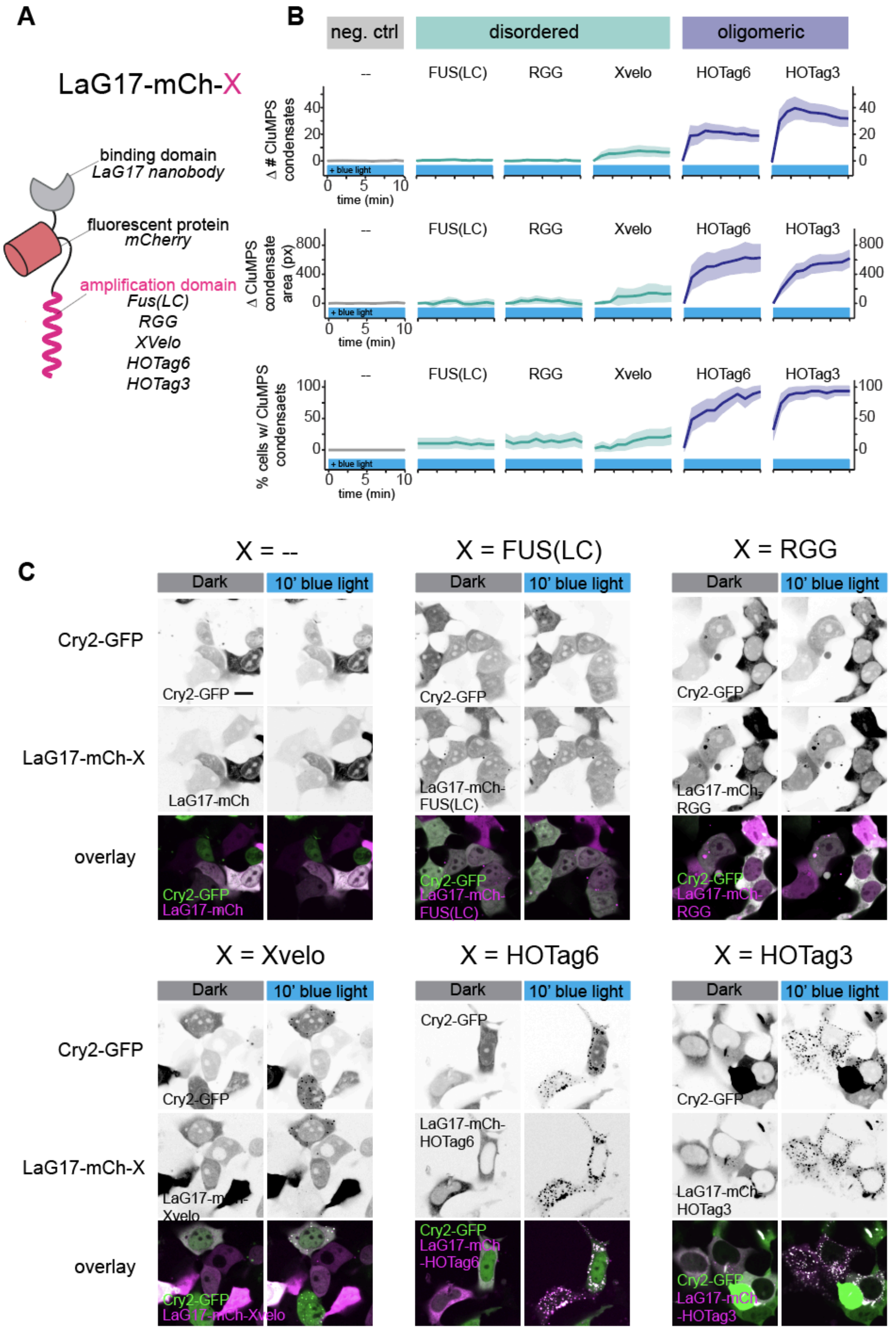


**Figure S1. Comparing candidate CluMPS amplification domains. (A)** Design of candidate CluMPS probes. CluMPS constructs comprise an N-terminal target-binding domain, a fluorescent protein, and a C-terminal “amplification domain” that can drive phase separation when clustered through binding to a clustered protein. Tested domains include the intrinsically disordered regions (IDRs) FUS(LC) (1-163), RGG, and Xvelo, and the homo-oligomeric tags HOTags 6 (tetrameric) and 3 (hexameric). A construct with no amplification domain was tested as a negative control. **(B)** Quantification of CluMPS condensate formation in cells cotransfected with Cry2-GFP and a CluMPS variant and stimulated with blue light over 10 minutes. Change in number of condensates per cell (top), change in area of clusters per cell (px, middle), and percentage of cells with clusters (defined as 5+ clusters AND 200+ pixels of cluster area (bottom) are shown for all constructs. Data represents mean, error bars = 95% CI for approximately 20-50 cells per group. **(C)** Representative images of all CluMPS variants co-expressed with Cry2-GFP in the dark and after 10 minutes of blue light stimulation. Scale bar = 20 μm.

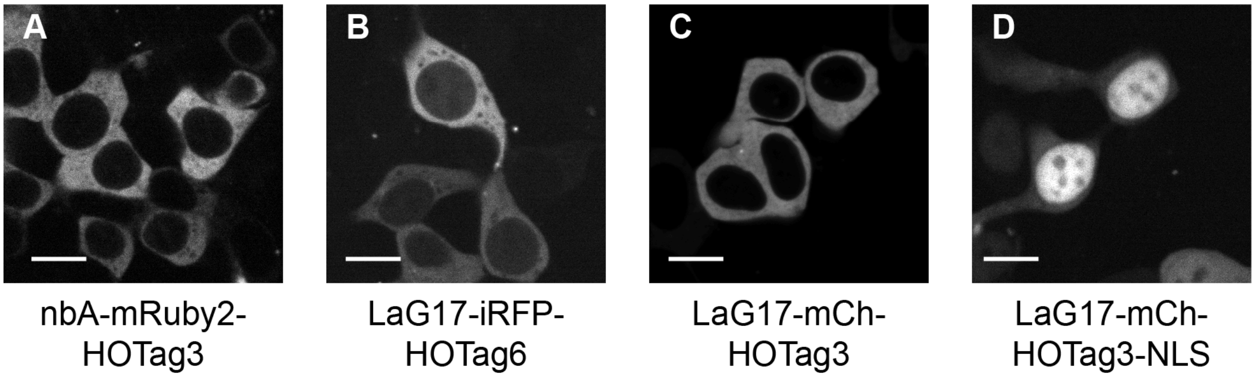


**Figure S2. CluMPS appears diffuse when transfected alone.** All CluMPS reporters generated appear diffuse when expressed alone in the absence of their target. Scale bars = 20 µm.

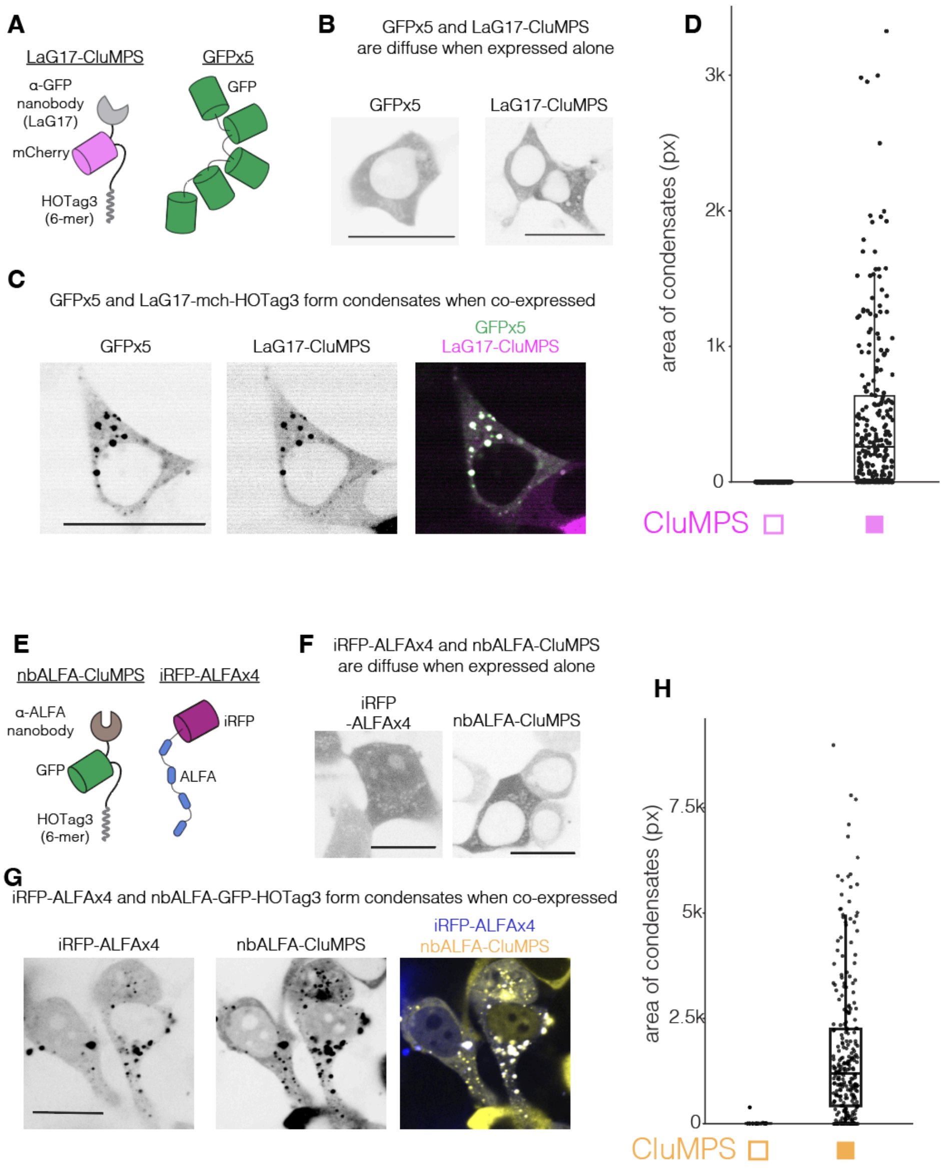


**Figure S3. CluMPS forms large condensates when coexpressed with oligomers of GFP or ALFAtag. (A)** LaG17 CluMPS (left) was tested against an oligomer of five tandem GFP molecules (GFPx5) fused onto a single peptide chain (right). **(B)** GFPx5 and LaG17-CluMPS appear diffuse when expressed separately. Scale bars = 20 μm. **(C)** GFPx5 and LaG17-CluMPS form large condensates containing both species when coexpressed. Scale bar = 20 μm. **(D)** Condensate area quantified for 338 cells containing only GFPx5 and 246 cells containing both GFPx5 and LaG17-CluMPS. Box-and-whisker plots represent median and quartiles and vertical lines extend from quartiles ± 1.5*(interquartile range). **(E)** nbALFA-CluMPS (left) was tested against an oligomer of four tandem ALFAtags fused onto a single peptide chain with miRFP670 (right), termed iRFP-ALFAx4. **(F)** iRFP-ALFAx4 and nbALFA-CluMPS appear diffuse when expressed separately. Scale bars = 20 μm. **(G)** iRFP-ALFAx4 and nbALFA-CluMPS form large condensates containing both species when coexpressed. Scale bar = 20 μm. **(H)** Condensate area quantified for 42 cells containing only iRFP-ALFAx4 and 267 cells containing both iRFP-ALFAx4 and nbALFA-CluMPS. Box-and-whisker plots represent median and quartiles and vertical lines extend from quartiles ± 1.5*(interquartile range). Scale bars = 20 μm.

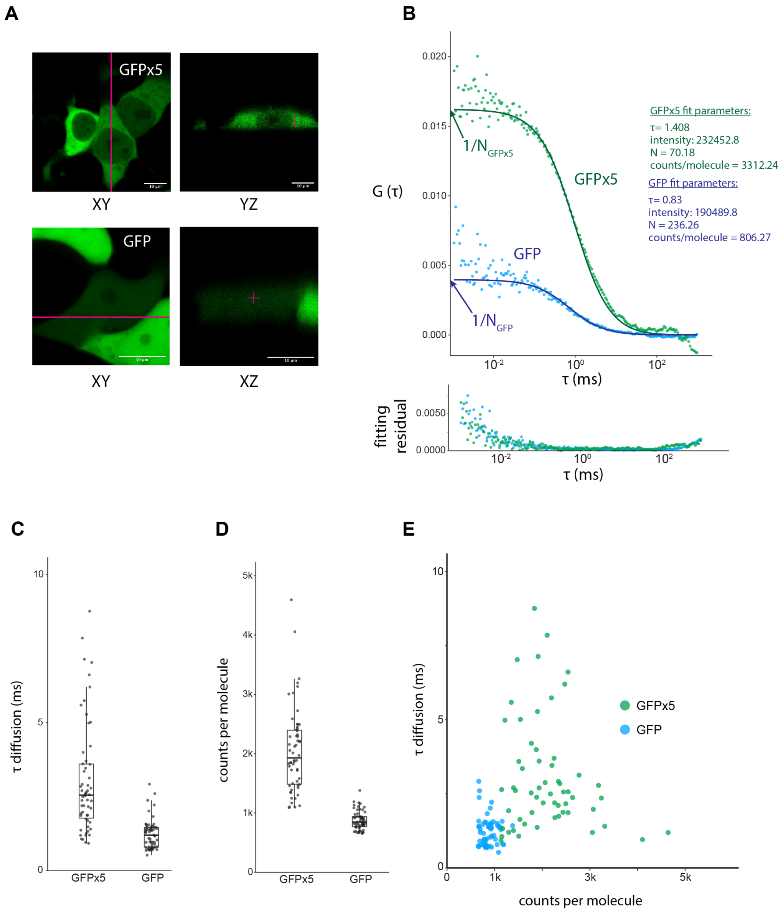


**Figure S4. Fluorescence correlation spectroscopy indicates that GFPx5 is not forming higher order aggregates in the absence of CluMPS**. We performed in-cell FCS to ensure that tandem-GFP constructs did not spontaneously aggregate in cells in the absence of CluMPS. **(A)** Representative fluorescence images of GFPx5 (top) and GFP (bottom), both of which appear diffuse via confocal imaging. Scale bars = 10 µm. **(B)** Representative FCS traces for GFPx5 (green) and GFP (blue). **(C)** Diffusion coefficients extracted from fitted FCS curves from cells transfected with GFP and GFPx5. On average, GFPx5 diffuses 1.7-fold slower than monomeric GFP. This closely matches the prediction of the Stokes-Einstein relation for the relative slowdown in diffusion for a molecule that is 5-fold larger. Stokes-Einstein states that a molecule's diffusion time is proportional to its hydrodynamic radius.

$$\tau_D \propto r$$

If we assume a spherical particle, then the hydrodynamic radius scales with the cube root of its molecular weight (MW).

$$r \propto \sqrt[3]{MW}$$

The ratio of diffusion times between GFPx5 and GFP will thus scale with the cube root of the ratio of their molecular weights

$$\frac{\tau_{D,\,GFPx5}}{\tau_{D,GFP}} \propto \frac{\sqrt[3]{MW_{GFPx5}}}{\sqrt[3]{MW_{GFP}}} = \sqrt[3]{\frac{MW_{GFPx5}}{MW_{GFP}}} = \sqrt[3]{5} \approx 1.7$$

**(D)** Molecular brightness (counts per molecule) of GFPx5 and GFP calculated as the average intensity divided by the number of molecules (N) obtained from fitted FCS curves. On average, GFPx5 was only 2-fold brighter, less than the potential 5-fold brightness increase, which may be accounted for by photobleaching, quenching, or incomplete maturation of all chromophores on a string. **(E)** Diffusion and counts per molecule of GFP or GFPx5 are largely uncorrelated, suggesting that neither species is forming higher order structures. Together, these data indicate that tandem GFP strings remain monomeric in cells and do not spontaneously self-associate.

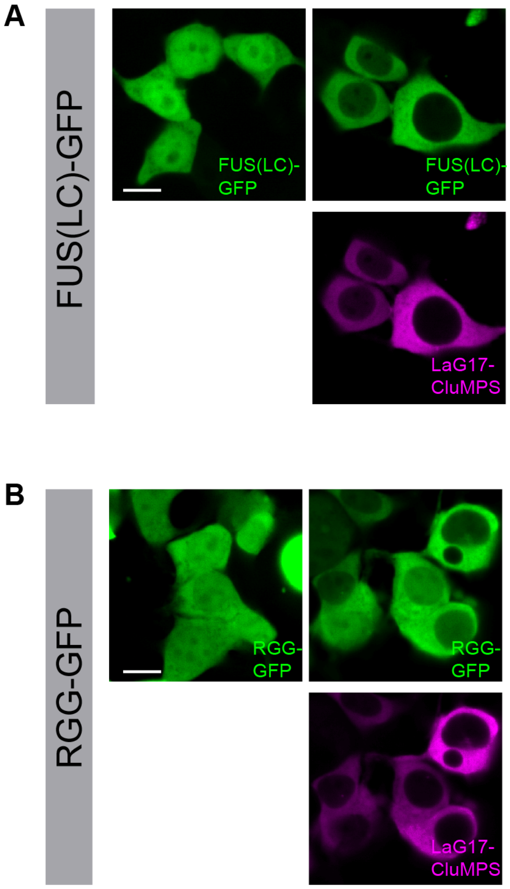


**Figure S5. LaG17-CluMPS does not trigger condensation of GFP fused to the IDRs FUS(LC) or RGG.** The lack of condensation of CluMPS when co-expressed with its target FUS(LC)-GFP **(A)** or RGG-GFP **(B)** suggests that CluMPS activation is specific to *bona fide* pre-existing clusters and does not give ‘false positive’ results by inducing clusters/condensates of a target where none existed. The altered nuclear localization of FUS(LC)-GFP and RGG-GFP in the presence of CluMPS confirms binding of CluMPS to its target despite lack of condensation. Scale bars = 20 µm.

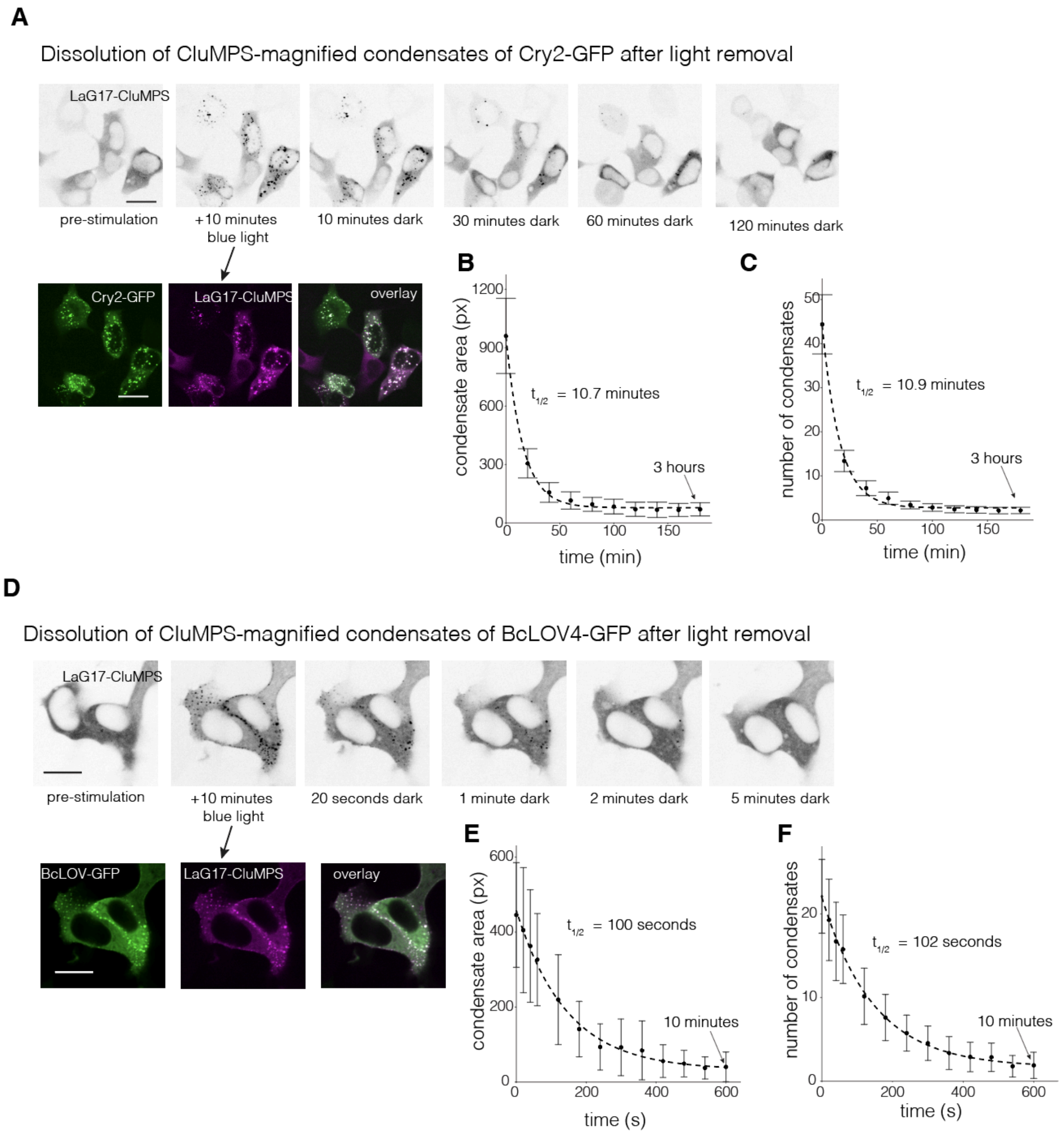


**Figure S6. Condensates of CluMPS and optogenetic clustering proteins dissolve when light stimulus is removed. (A)** Representative images of condensates of LaG17-CluMPS and Cry2-GFP forming during 10 minutes of blue light stimulation and dissolving over ~2 hours after removal of blue light. Scale bars = 20 µm. Decay kinetics of condensate area **(B)** and number **(C)** after light removal. **(D)** Representative images of condensates of LaG17-CluMPS and BcLOV4-GFP forming during 10 minutes of blue light stimulation and dissolving over ~5 minutes after light. Scale bars = 20 µm. Decay kinetics of condensate area **(E)** and number **(F)** after light removal.

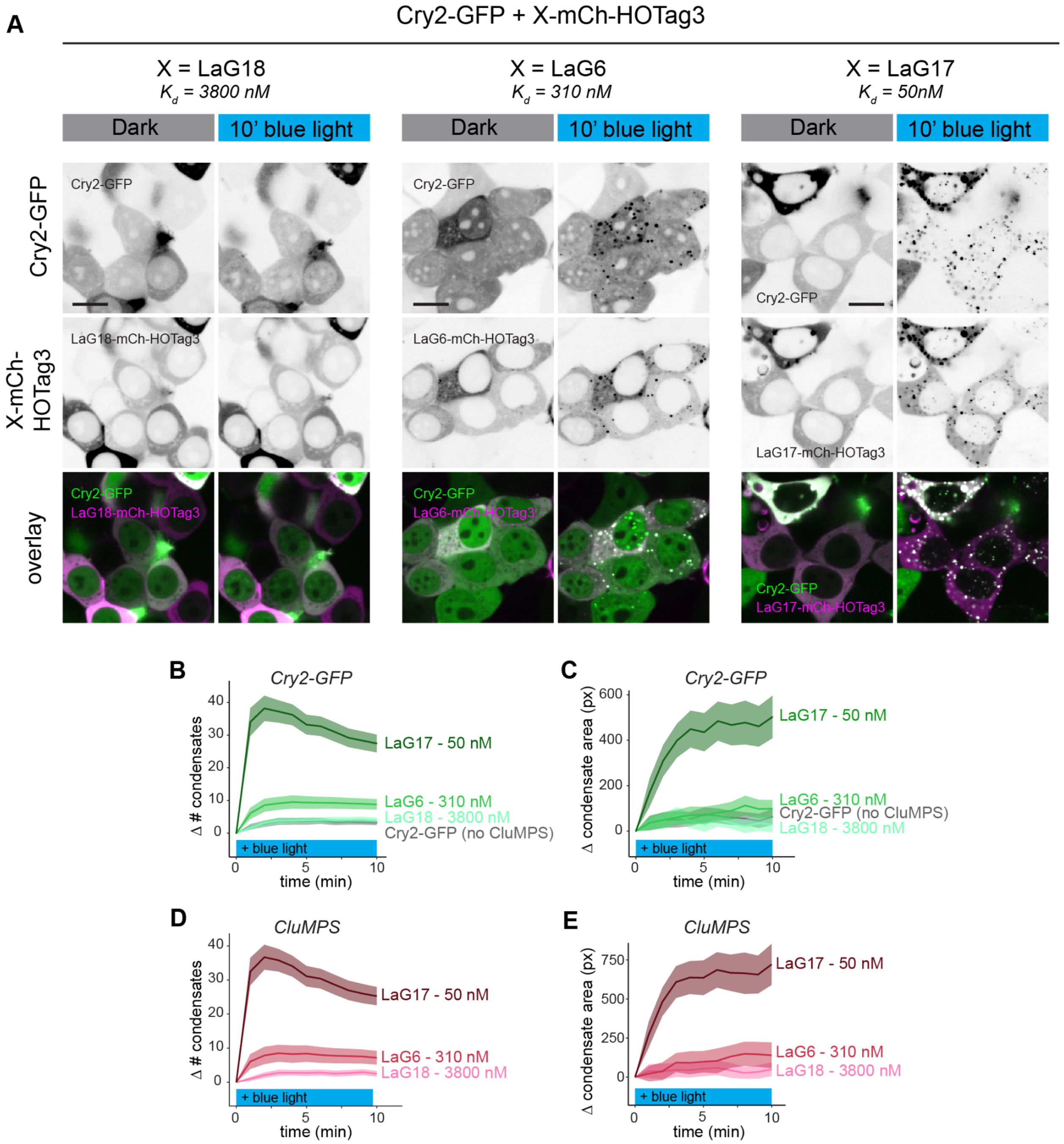


**Figure S7. CluMPS activity depends on the affinity of the target binding domain.** To test the effect of binding domain affinity on CluMPS activation, we co-expressed Cry2-GFP with CluMPS variants with binding domains with differing affinities for GFP. Binding domains used were LaG18 ($K_d$ = 3800 nM), LaG6($K_d$ = 310 nM) and LaG17 ($K_d$ = 50 nM). **(A)** Representative images of condensation of both Cry2-GFP and CluMPS both before and after 10 min of stimulation with blue light. Scale bars = 20 µm. Quantification of condensate number (**B,D**) and area (**C,E**) in both the GFP (**B,C)** and miRFP (**D,E**) shows that stronger binding between CluMPS and target yields stronger condensation. Each trace represents mean, ribbons = 95% CI for approximately 100-200 cells per group.

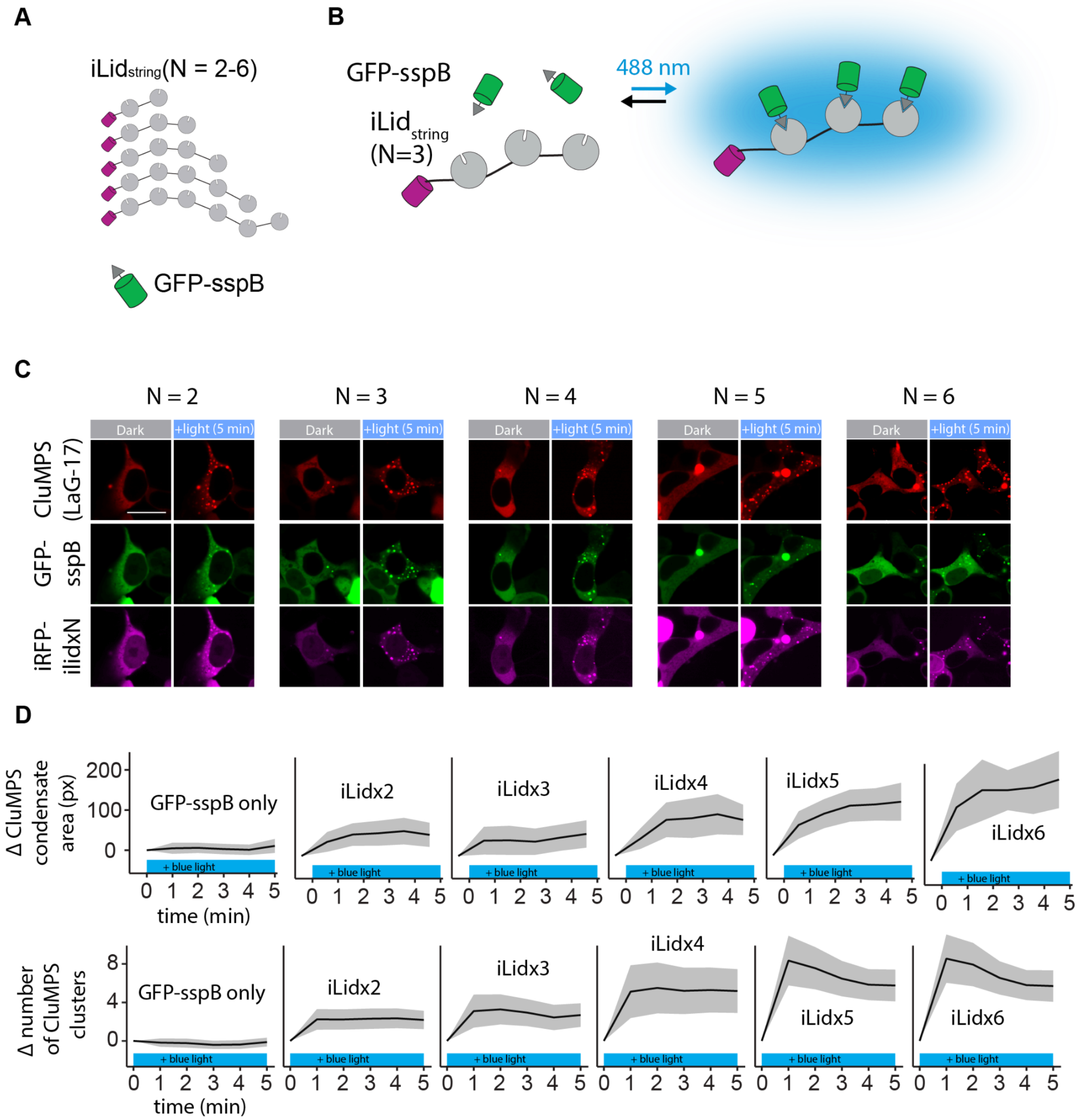


**Figure S8. iLid$_{strings}$ and GFP-sspB form defined multimers upon blue light stimulation, providing a platform for characterizing CluMPS sensitivity to small multimers. (A)** Multiple iLids on a single peptide, termed iLid$_{strings}$, span between two and six iLids, determining the number of GFP-sspB monomers which are recruited to form a small oligomer. (**B)** GFP-sspB is recruited to iLid$_{string}$ upon blue light exposure. (**C)** Representative images of CluMPS condensation, which can be observed for all iLid$_{string}$ (length 2-6) and GFP-sspB under ideal conditions. Scale bar = 20 µm. (**D)** Change in CluMPS condensate area (pixels, top) and number of CluMPS condensates after blue light stimulation for GFP-sspB only, and GFP-sspB expressed with iLid$_{string}$ (length 2-6). Data represents mean traces of 150-200 cells per condition, ribbon shows 95% CI.

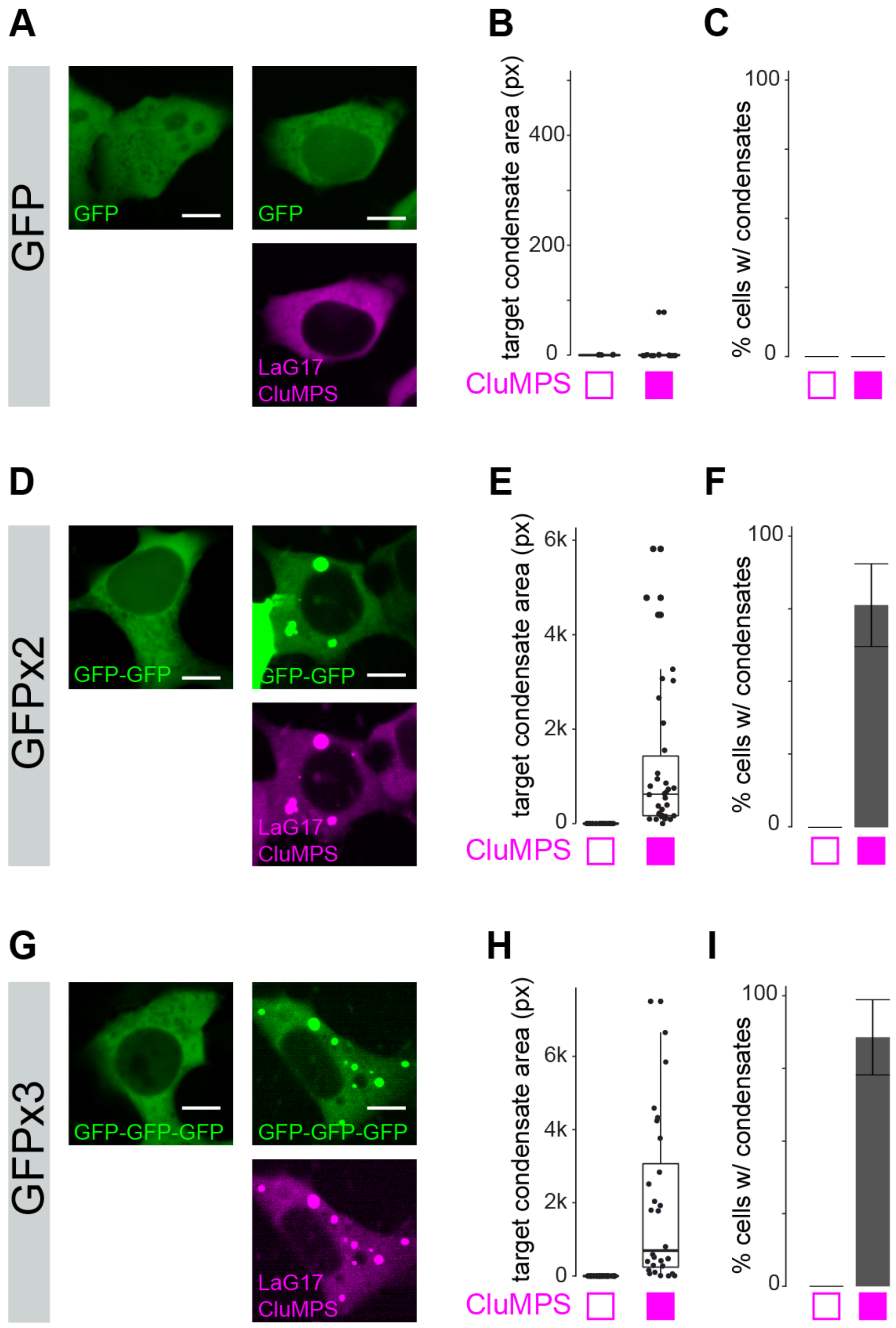


**Figure S9. Tandem GFP trimer and dimer, but not GFP monomer, triggers CluMPS condensation.** Representative images of GFP(**A**), GFP-GFP (**D**), or GFP-GFP-GFP (**G**) transfected in the presence or absence of LaG17-mCh-HOTag3 (LaG17-CluMPS). Scale bars = 20 µm. Quantification of GFP condensate area (**B,E,H**) and % of cells with GFP condensates (**C,F,I**) shows CluMPS-induced condensation with the multimeric GFPs, but not with monomeric GFP. Box-and-whisker plots in (**B,E,H**) represent median and quartiles from approximately 20-30 cells per group. Vertical lines extend from quartiles ± 1.5*(interquartile range). Bar plots (**C,F,I**) represent frequency. Error bars = 95% confidence interval of approximately 20-30 cells per group.

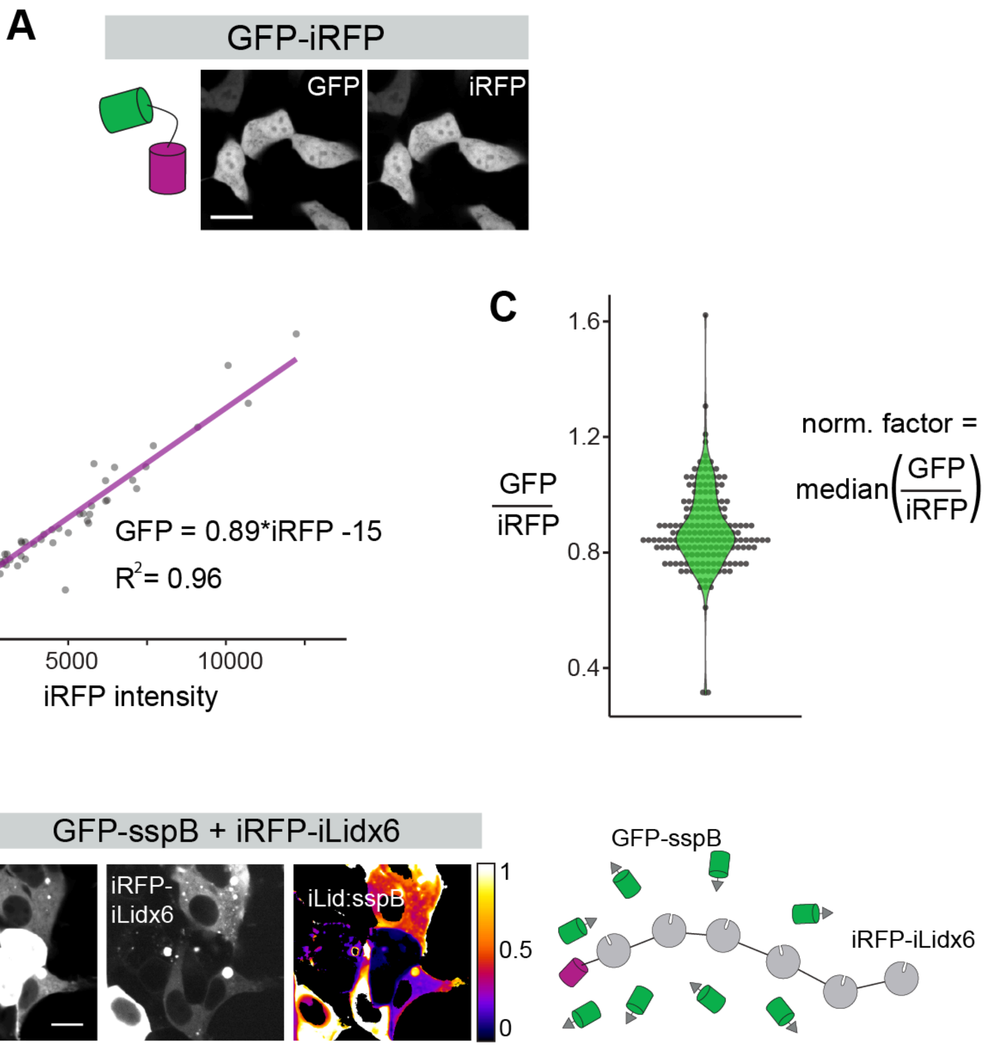


$$\text{iLid:sspB} = \frac{\text{iRFP-iLidx6 intensity}}{\text{GFP-sspB intensity}} * \text{norm. factor} * \text{6 iLid/iRFP}$$

$$\text{\% GFP clustered} = \min(100,\ 100*\text{iLid:sspB})$$

**Figure S10. Using GFP-iRFP as a control for determining stoichiometric ratio of GFP-sspB and iRFP-iLid$_{string}$.** The relative stoichiometries of GFP-sspB and iRFP-iLid$_{string}$ were determined by separately imaging a direct fusion of GFP-iRFP, which expressed both fluorophores in a 1:1 ratio. **(A)** Representative images of EGFP-miRFP670 (GFP-iRFP) in HEK 293T. Scale bar = 20 µm. **(B)** Plot of GFP intensity vs iRFP intensity shows a linear relationship ($R^2$ = 0.96) between expression of the two fluorophores. Note that linear agreement was obtained only when supplementing cells with exogenous iRFP cofactor biliverdin (see **Methods** for details). **(C)** Ratio of GFP to iRFP intensity for individual cells with overlaid violin plot. The normalization factor used in subsequent calculations is the median GFP:iRFP ratio. **(D)** Representative image of GFP-sspB (left), iRFP-iLidx6 (middle), and iLid:sspB ratio pseudo-image (right), and corresponding cartoon showing GFP-sspB and iRFP-iLidx6. Pseudo-image generated according to the equation shown: iRFP intensities are divided by GFP intensities, normalized with the GFP-iRFP normalization factor from (**C**), and multiplied by the number of iLids per iRFP on each string (in this case, 6). The percentage of GFP clustered is the percentage of GFP-sspB molecules that have iLid sites to bind on the multimer and is calculated by multiplying the iLid:sspB ratio by 100 (calculated values over 100 are considered 100% clustered). Note that although the pseudoimage shows pixel-level iLid:sspB values, the iLid:sspB ratio and corresponding % target clustered in the manuscript are cell-level metrics that were calculated from the median values of the GFP and iRFP channels. Scale bar = 20 µm.

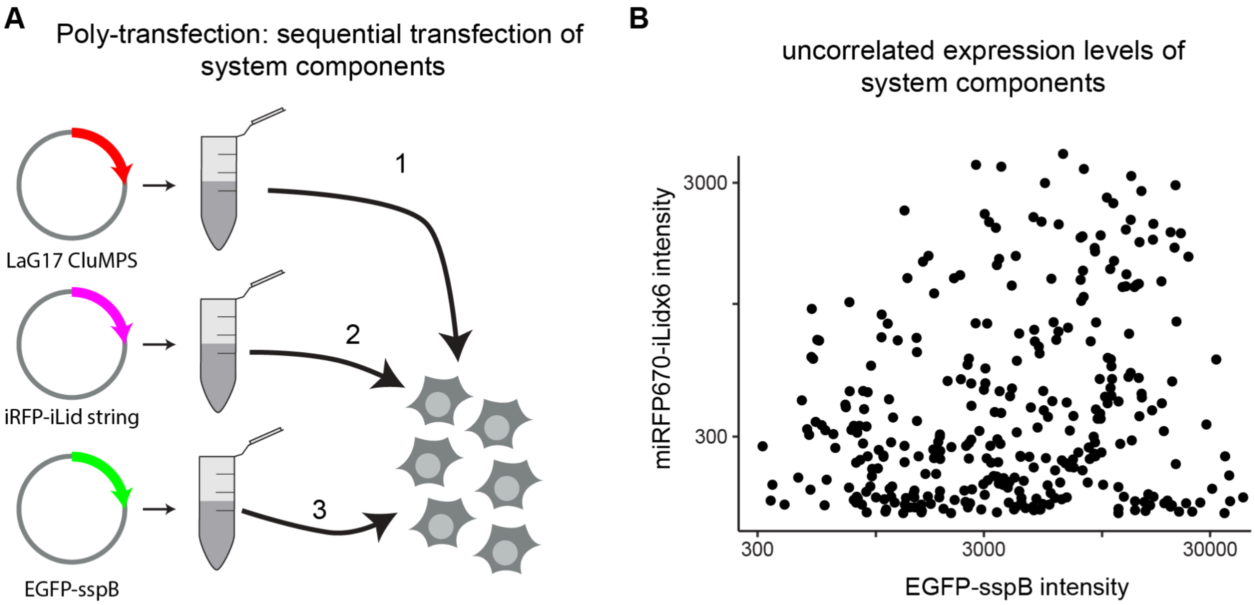


**Figure S11. Poly-transfection allows transfection of multiple components with low expression correlation and allows sampling of a wide range of iLid:sspB ratios. (A)** Cartoon depicting poly-transfection[71]. Plasmids were separately mixed with transfection reagent and sequentially added to cells so that their subsequent expression would be largely uncorrelated. **(B)** Plot of miRFP670 intensity vs EGFP intensity after poly-transfection shows no correlation between iRFP and GFP levels.

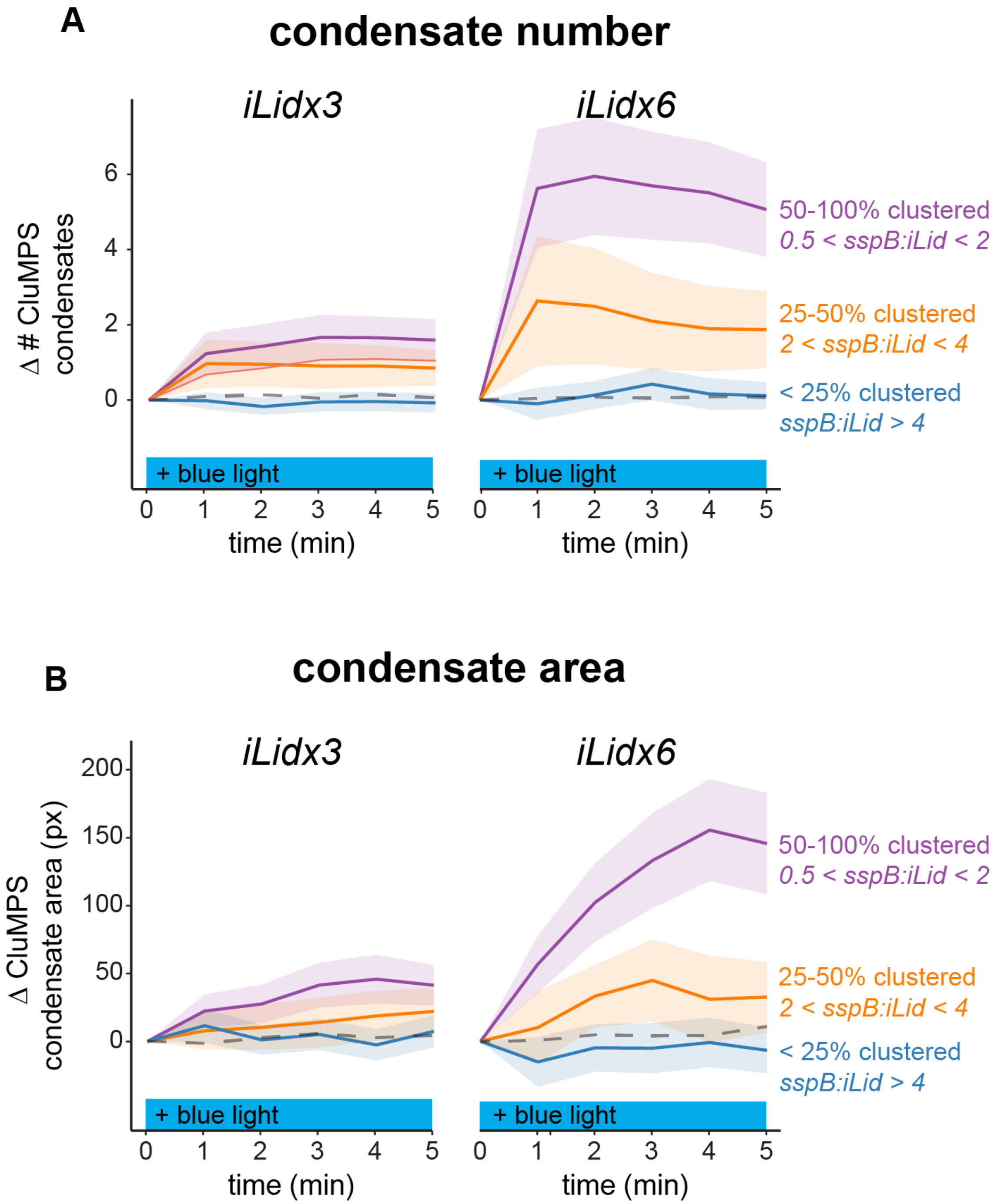


**Figure S12. CluMPS activation depends on both cluster size and the fraction of target that exists in a cluster.** Condensation was quantified for HEK 293T cells transfected with LaG17-CluMPS, GFP-sspB, and iLid strings (of size 3 or 6). The number (**A**) and size (**B**) of condensates was measured during 10 min of blue light stimulation. Condensation was stronger for cells co-transfected with iLidx6, demonstrating how larger cluster sizes promote stronger CluMPS activation. However, for both target cluster sizes, condensation also depended on the estimated number of sspB monomers in a cluster, which was estimated based on the stoichiometry of sspB and iLid in single cells. Dotted gray line shows change in number of clusters after blue light stimulation in cells from the same pool that were transfected with CluMPS and GFP-sspB but not with iLid string, and therefore cannot form light-induced clusters. Each trace represents mean, ribbons = 95% CI for approximately 50-200 cells per group.

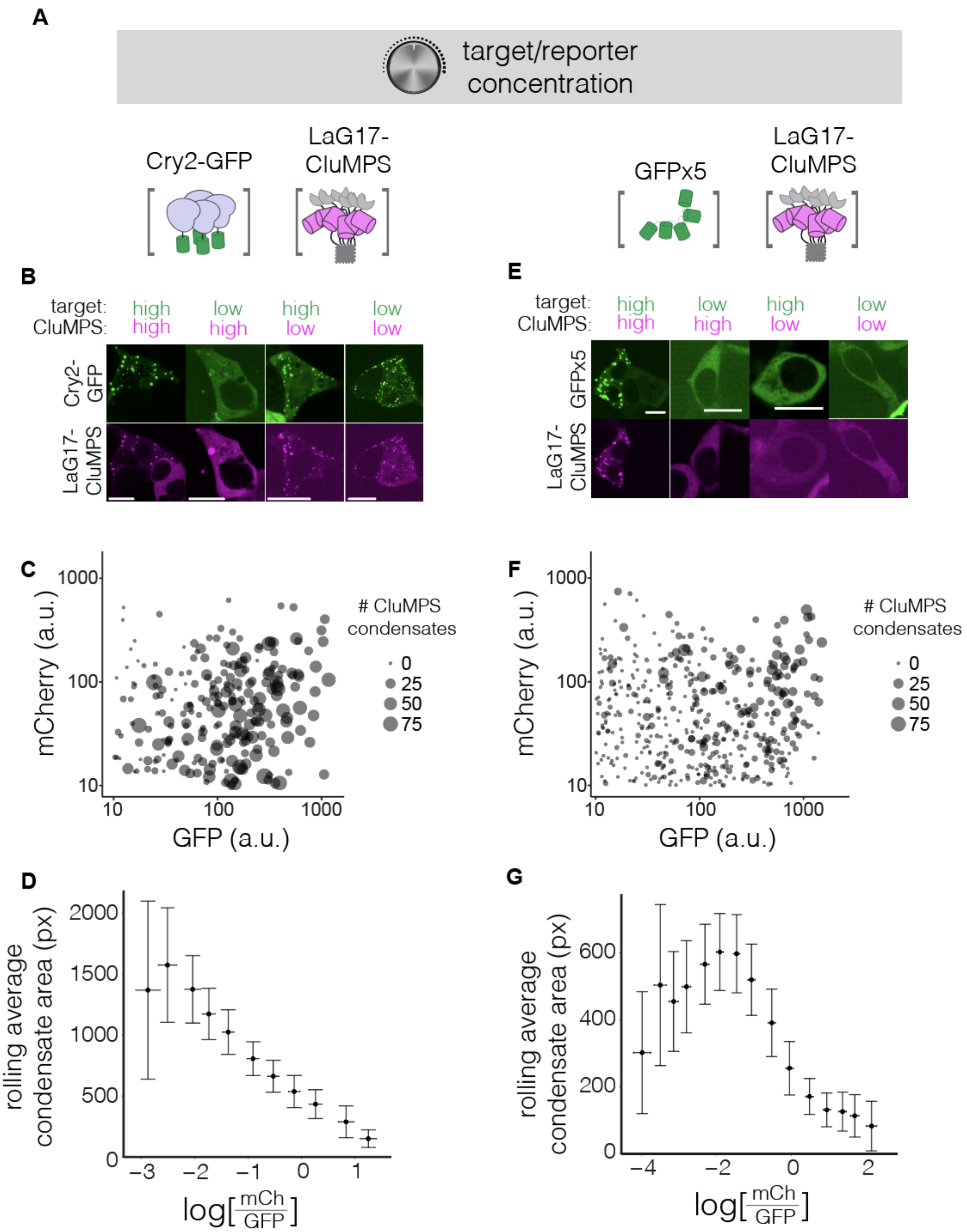


**Figure S13. Condensation of LaG17-CluMPS and target depends on the ratio of concentrations of the two components. (A)** Cry2 and GFPx5 were used as two model target clusters measure the effect of CluMPS and target concentration on CluMPS activation. **(B)** Representative images of HEK 293T cells expressing LaG17-CluMPS and Cry2-GFP. Condensation was observed in all expression regimes except when Cry2 was low and CluMPS was high. Scale bar = 20 µm. **(C)** Scatter plot of condensation as a function of concentration of the two components. Each dot represents a cell, and dot size represents number of condensates in that cell. **(D)** Condensation decreased with increasing CluMPS:Cry2 ratio. **(E)** Representative images of HEK 293T cells expressing LaG17-CluMPS and GFPx5. Condensation was observed only when both CluMPS and target were sufficiently high. Scale bar = 20 µm. **(F)** Scatter plot of condensation as a function of concentration of the two components. Each dot represents a cell, and dot size represents the number of condensates in that cell. **(G)** Condensation was strongest at an intermediate CluMPS:GFPx5 ratio and decreased with increased expression bias towards one component.

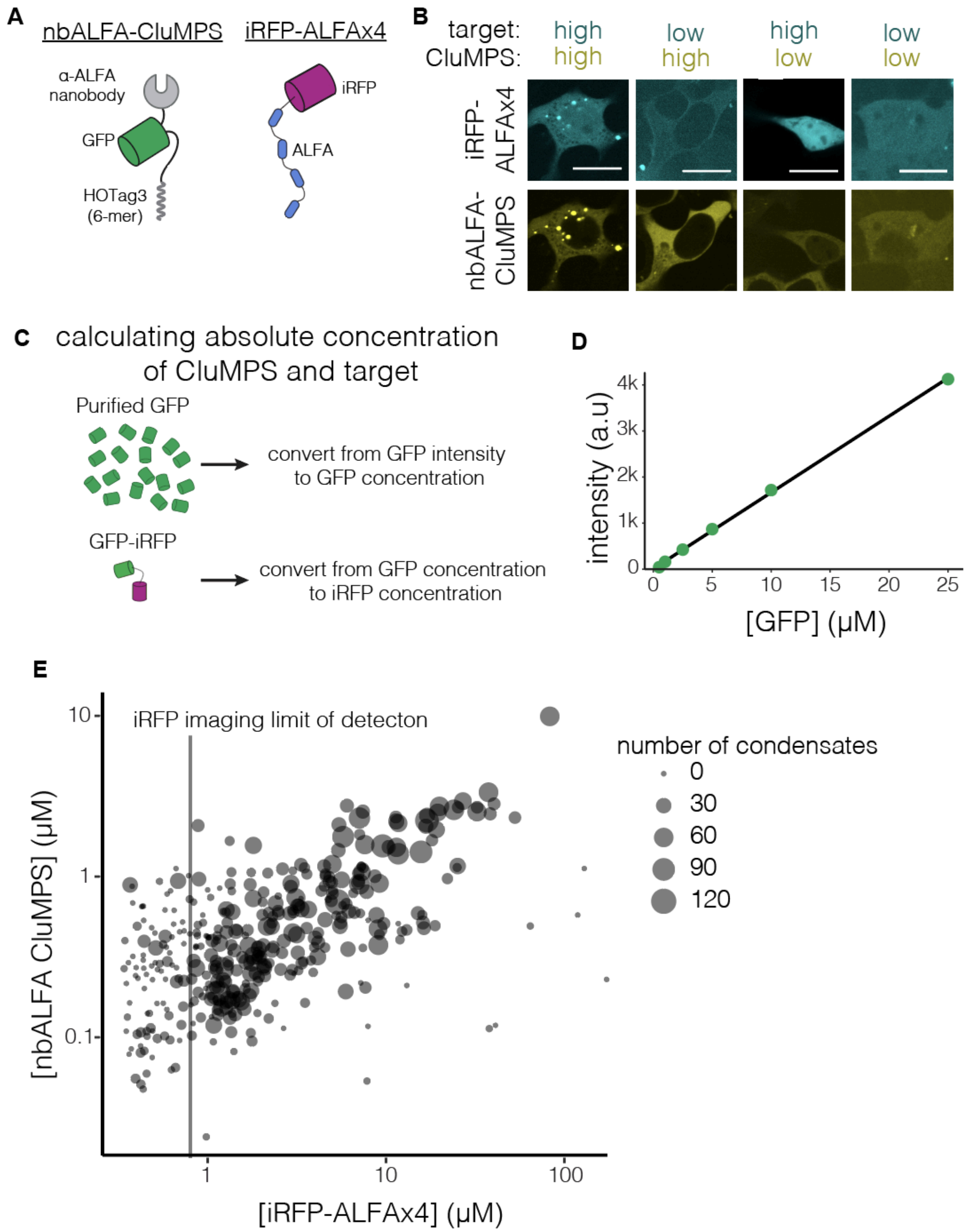


**Figure S14. Obtaining an estimate for the absolute concentration regime required for CluMPS activation of small clusters. (A)** A test target of four tandem ALFAtags fused to miRFP670 can be targeted with CluMPS harboring an ALFA-binding nanobody (nbALFA) as a binding domain. **(B)** Representative images of CluMPS condensation as a function of CluMPS and target expression. Condensation required above-threshold expression for both components, similar to results seen with GFPx5. Scale bars = 20 μm. **(C)** Absolute concentration of CluMPS and target in single cells was estimated by using purified GFP of known concentration to convert from image intensity to GFP concentration. This fluorescence was then compared to cells transfected with GFP-iRFP, allowing calibration of iRFP intensity to iRFP concentration (see **Methods**). **(D)** Dilutions of purified GFP were imaged and the intensity was averaged over the image. Concentration of GFP and image intensity are linearly related ($R^2$ = 0.9994). **(E)** Absolute concentration of nbALFA-CluMPS and iRFP-ALFAx4, with point size scaled to number of condensates. Higher concentration and proper ratio of both components improves clustering. Vertical line at ~0.8 μM shows limit of detection for iRFP-ALFAx4 (cells to the left of this line did not have sufficient transfection of iRFP-ALFAx4 to be detectable via imaging).

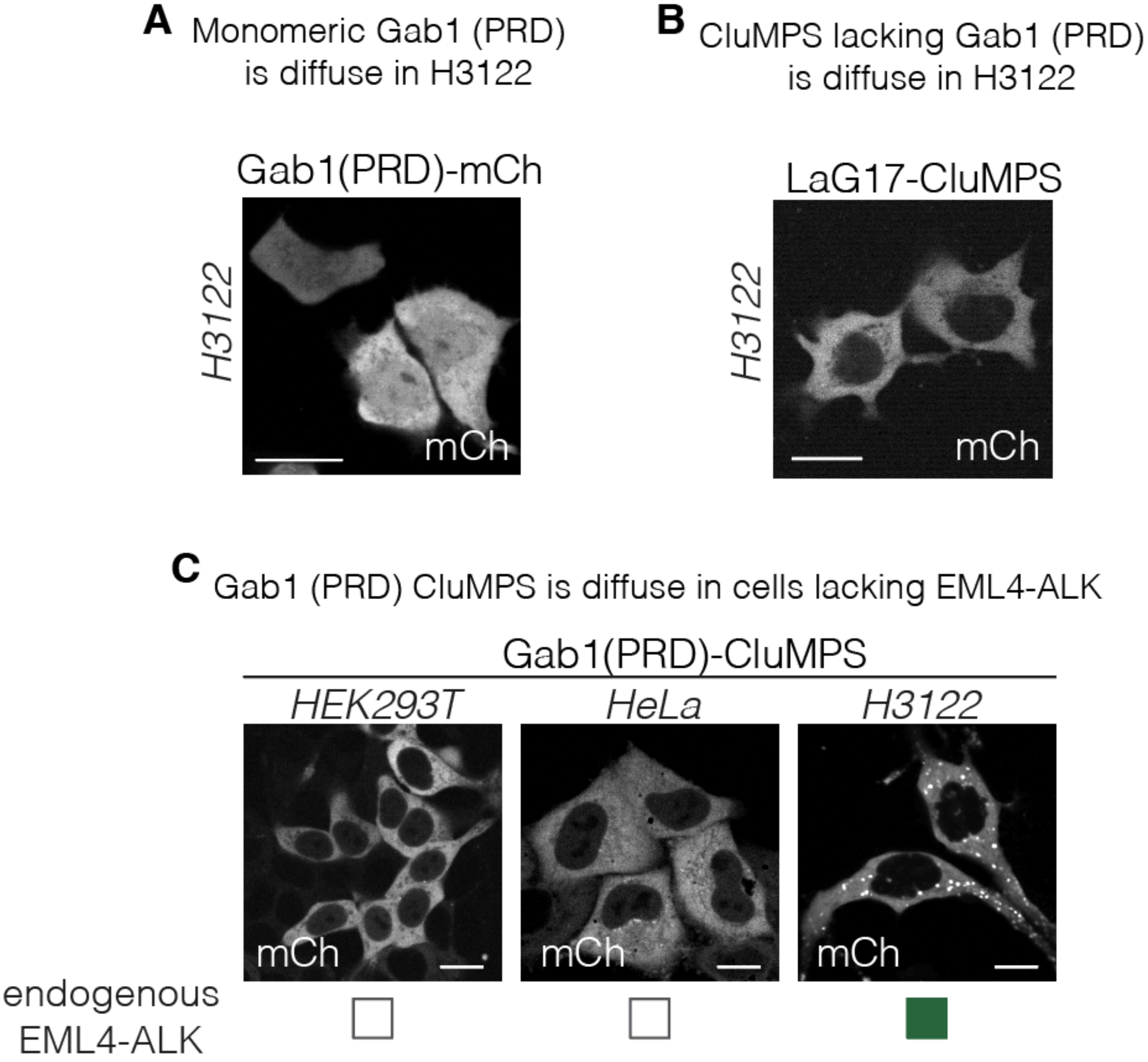


**Figure S15. Gab1(PRD)-CluMPS is diffuse in cells that lack the EML4-ALK oncogene but is clustered in cells that express EML4-ALK. (A)** Monomeric Gab1 (PRD)-mCh is diffuse in H3122 cells. **(B)** A CluMPS probe that lacks the Gab1 (PRD) domain is diffuse in H3122 cells. **(C)** Gab1(PRD)-CluMPS expressed in various cell types. (left) Gab1(PRD)-CluMPS is diffuse in HEK293T cells. (center) Gab1(PRD)-CluMPS is diffuse in HeLa cells. (right) Gab1(PRD)-CluMPS forms condensates in H3122 cells, which express endogenous EML4-ALK. The expression levels depicted are roughly equivalent between the different cell lines. All scale bars = 20 µm.

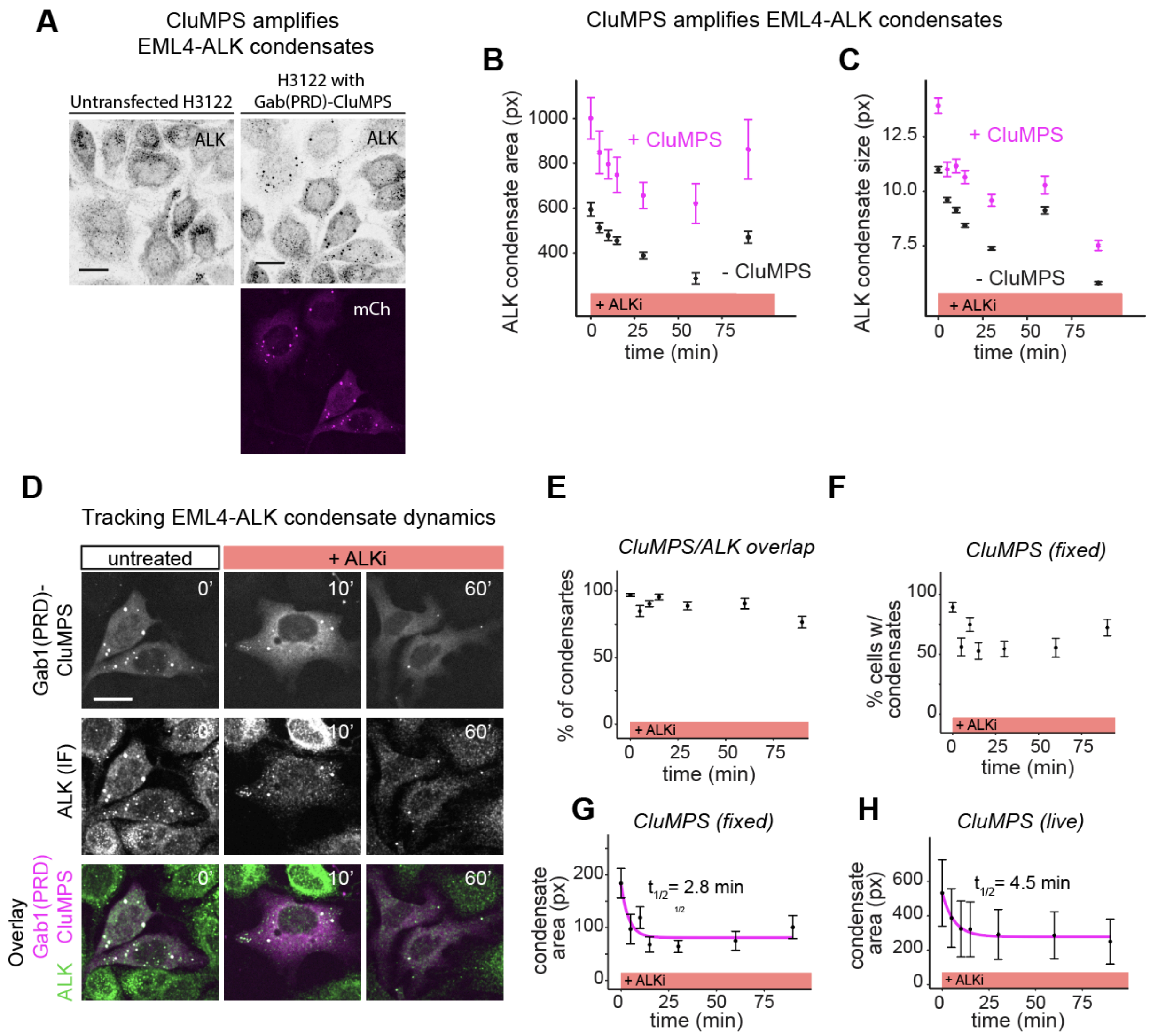


**Figure S16. Gab1(PRD)-CluMPS amplifies EML4-ALK condensates and visualizes their dynamics in H3122 cancer cells. (A)** Immunofluorescence images of H3122 cells stained of ALK with and without Gab1(PRD)-CluMPS. Scale bars = 20 µm. Comparison of ALK condensate area per cell **(B)** and ALK condensate mean size **(C)** in cells with and without CluMPS demonstrates that CluMPS amplifies EML4-ALK condensates. Data in **(B)** represents mean, error bars = 95% CI of between 150-250 cells with CluMPS and 1200-1800 cells without CluMPS. Data in **(C)** represent mean, error bars = 95% CI for approximately 40,000-110,000 clusters per group in cells without CluMPS, and 9000-18000 clusters per group in cells transfected with CluMPS. **(D)** Representative immunofluorescence images of H3122 cells expressing Gab1(PRD)-CluMPS before and during treatment with 1 µM crizotinib. Scale bar = 20 µm. **(E)** Percentage of all CluMPS condensates that overlapped with ALK (defined as having >10% overlap of pixels identified as both a CluMPS and ALK condensate) at each time point of drug treatment. Data represents percentage, error bars = 95% CI of between 200-1000 condensates per time point. **(F)** Percentage of cells at each time point containing CluMPS condensates (defined as having >1 cluster and >25 total pixels of CluMPS condensate area). Data represents percentage, error bars = 95% CI of between 150-250 cells per time point. **(G)** Area of CluMPS condensates (pixels per cell) in fixed cells after different durations of drug treatment. Exponential decay fit overlayed in magenta. Data represents mean, error bars = 95% CI of between 150-250 cells per time point. **(H)** Area of CluMPS condensates (pixels per cell) from live cell imaging of H3122 cells treated with 1 µM crizotinib, reproduced from **Figure 5E** and presented here for side-by-side comparison with fixed cells. Exponential decay fit overlayed in magenta. Data represents mean, error bars = 95% CI of 31 cell traces.

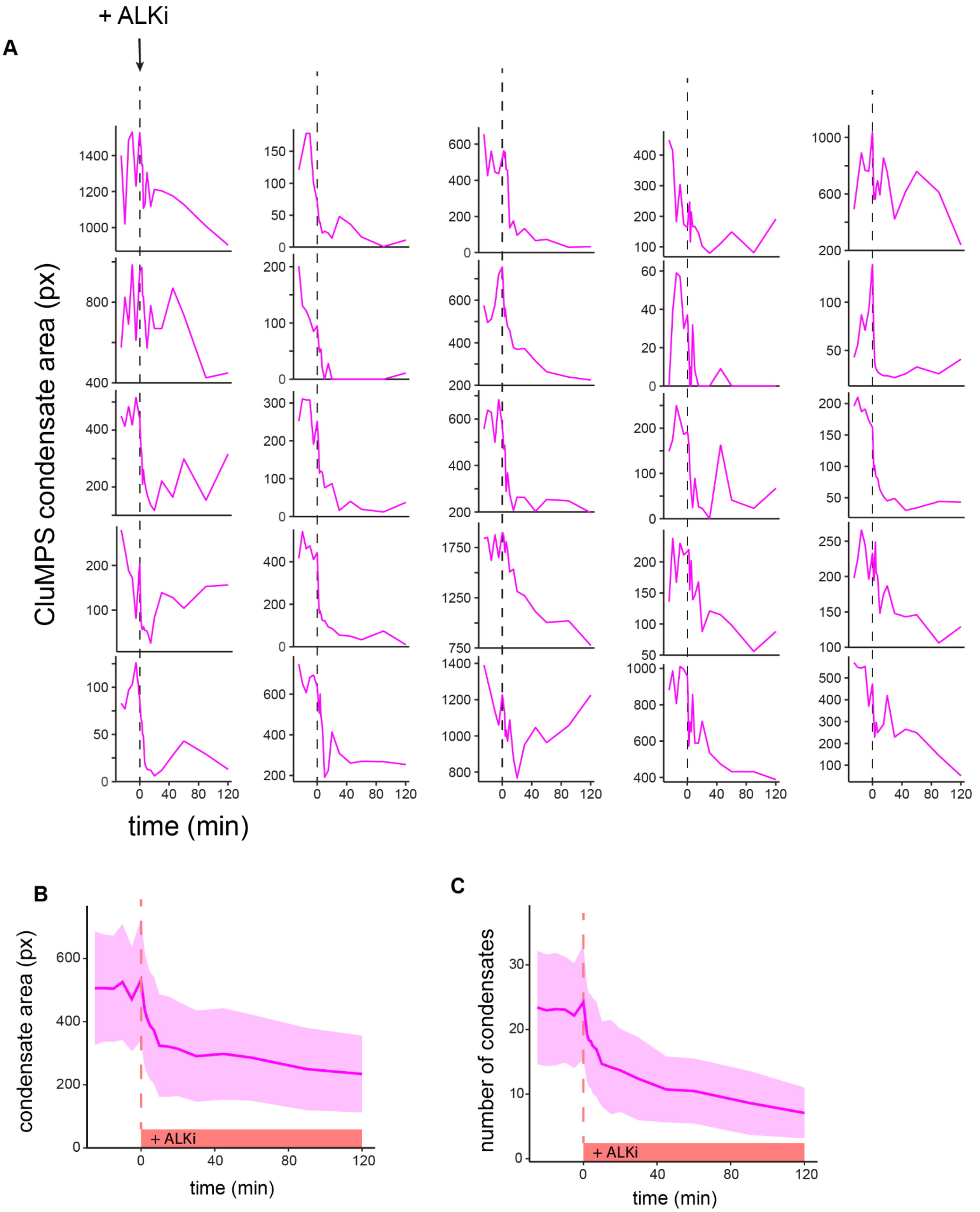


**Figure S17. Single-cell visualization of Gab1(PRD)-CluMPS dynamics through live-cell imaging of H3122 cells. (A)** Single-cell traces of multiplexed CluMPS in response to ALK inhibitor (1 μM crizotinib). Dotted line indicates when drug was added. **(B,C)** Average CluMPS condensate area and number in H3122 cells after crizotinib treatment. **(B)** is reproduced from **Figure 5E**. Traces represents mean, ribbon = 95% CI of 31 cell traces.

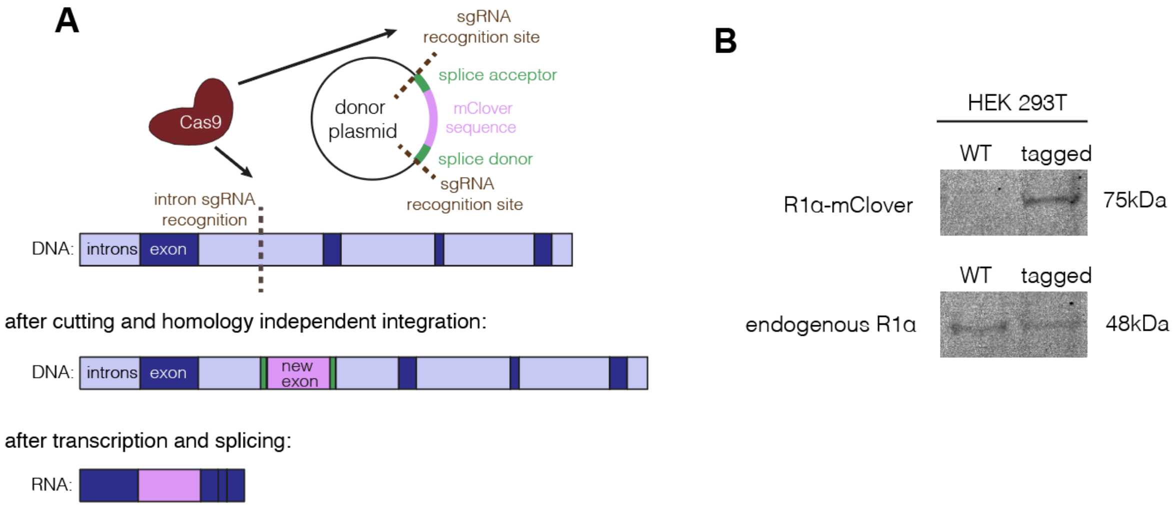


**Figure S18 - Figure showing strategy and confirmation of successful tagging of R1α**
**(A)** Schematic depicting method of tagging PKA R1α with mClover3, adapted from Serebrenik et al. **(B)** Western blot showing endogenous R1a band present in wild type and in tagged cells, while mClover-tagged R1α appears in second band at 75 kDa not seen in wild type cells.

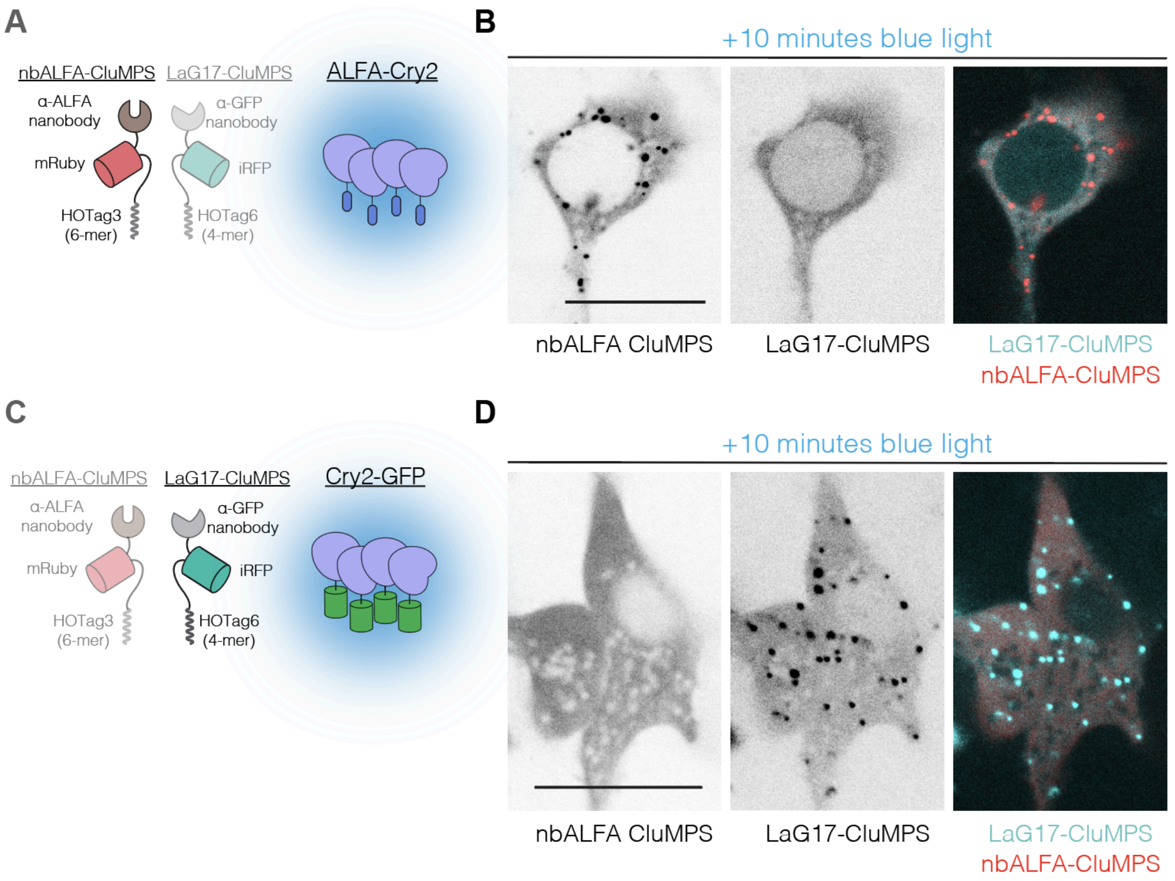


**Figure S19. LaG17-iRFP-HOTag6 and nbALFA-mRuby-HOTag3 are orthogonal CluMPS probes that do not visibly interact with each other. (A)** Cells were transfected with LaG17-CluMPS, nbALFA-CluMPS, and ALFA-Cry2. **(B)** After blue light induced clustering of Cry2, condensates of ALFA-Cry2 and nbALFA-CluMPS form, while LaG17-CluMPS remains diffuse. **(C)** Cells were transfected with LaG17-CluMPS, nbALFA-CluMPS, and Cry2-GFP. **(D)** After blue light induced clustering of Cry2, condensates of Cry2-GFP and LaG17-CluMPS form, while nbALFA-CluMPS remains diffuse. Scale bars = 20 μm.

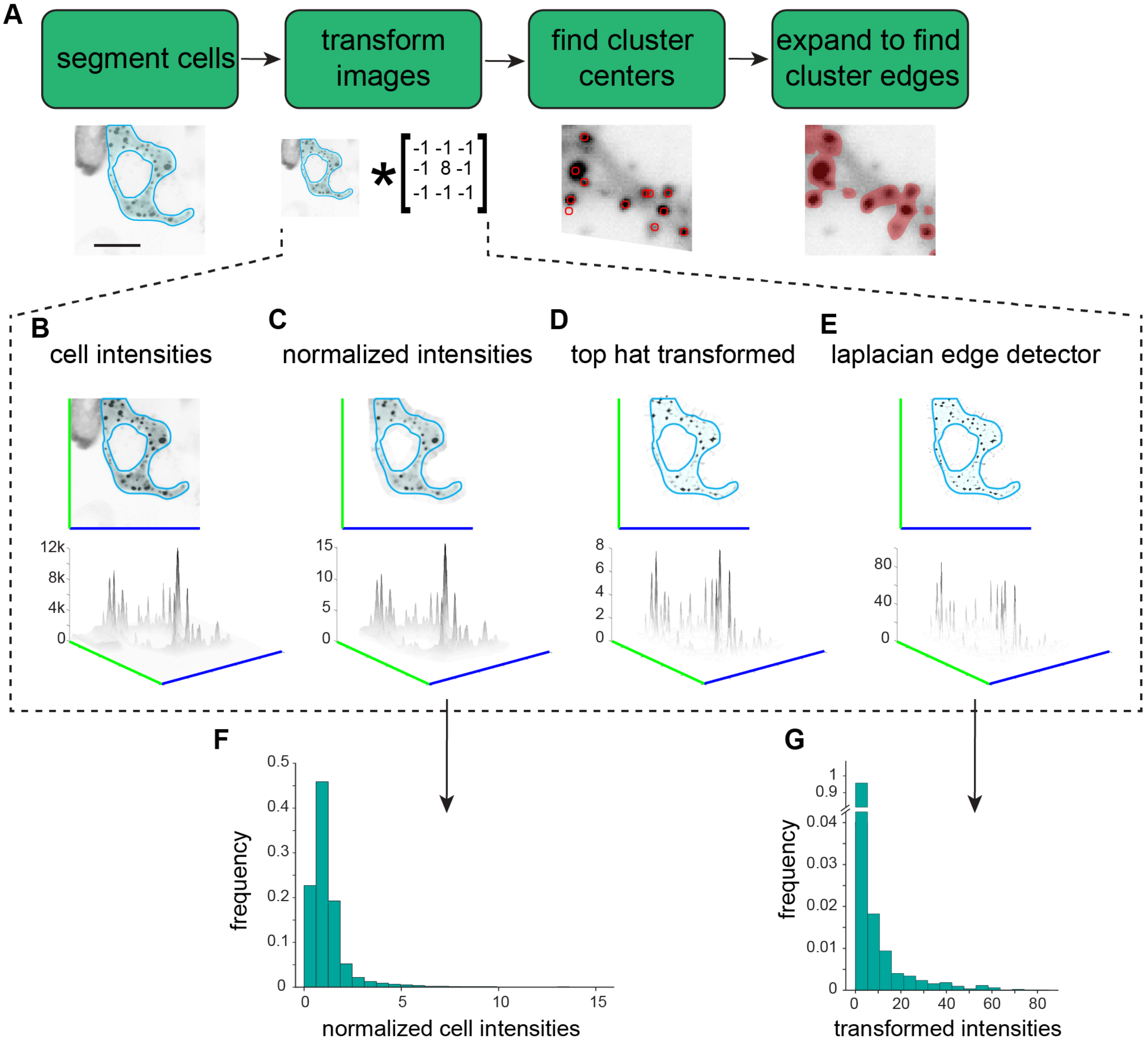


**Figure S20. Fitting clusters in single cells with a custom MATLAB script: flow chart and image transformation**. A custom MATLAB algorithm was developed to segment cells and identify clusters (see **Methods** and associated code files for more details). **(A)** Flow chart showing basic steps of cluster fitting. First, single cells were segmented, and cell intensities were transformed to normalize them, suppress background, and enhance clusters (see **C-E** below). This transformed image was used to identify points at the center of clusters, and finally a stepwise algorithm expanded from each cluster center to determine the pixels within each cluster. Scale bar = 20 µm. Further details on image transformation: After segmentation **(B)**, image intensities were normalized within a cell mask by subtracting background and dividing by the median. **(C)** A Top hat transform suppressed all features on longer length scales than desired, effectively removing background fluorescence within the cell (non-clusters go to zero). **(D)** Next, a laplacian edge detector transformation enhanced contrast and further amplified the intensities of areas of high contrast (clusters). **(E)** A histogram of intensities after the normalization stage shows most pixels around 1, and the brightest pixels between 3 and 15. **(F)** By contrast, a histogram of final transformed values shows >95% of pixels ~0, and the brightest pixels are 15-80, thus enhancing cluster contrast. In sum, this procedure successfully suppressed background fluorescence while enhancing clusters, enabling robust, semi-automated detection of fluorescent clusters.

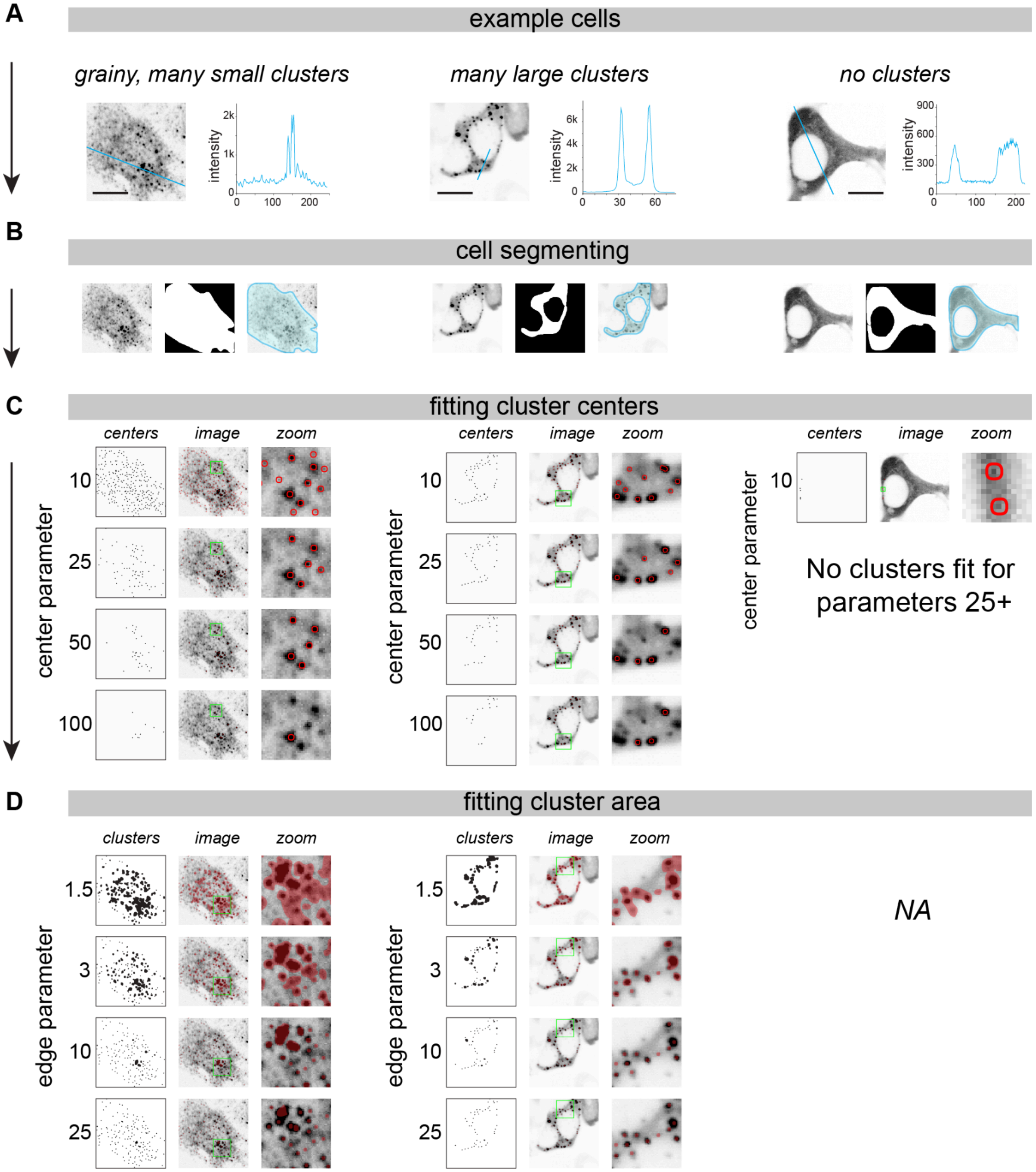


**Figure S21. Fitting clusters in single cells with a custom MATLAB script: two parameters determine how clusters are fit. (A)** Cluster fitting is depicted for three classes of clustered cells. Left, a cell with grainy fluorescence and small clusters. Middle, a cell with many clearly-defined clusters. Right, a cell with no clusters. Scale bars = 20 µm. **(B)** Cell masks are generated from initial cell segmentation (see **Methods** for more details). **(C)** The first fitting parameter (center parameter) determines how strictly cluster centers are fit (see **Methods**), with higher values being more strict and resulting in fewer identified clusters. **(D)** The second (edge) parameter determines how strictly the edges of clusters are selected, with higher values being more strict and resulting in smaller cluster areas.
